# miR319 promotes *de novo* shoot regeneration by repressing *LsTCP4* in lettuce

**DOI:** 10.64898/2026.07.08.737254

**Authors:** Tao Jiang, Sameena Ejaz Tanwir, Anandi Karn, Fangchen Liu, Heqiang Huo

**Affiliations:** Crop Transformation Center, Department of Horticulture Sciences, IFAS, University of Florida, Gainesville, FL 32611, USA; Mid-Florida Research & Education Center, IFAS, University of Florida, Apopka, FL 32703, USA

**Author notes:** First author: Tao Jiang. Correspondence: Heqiang Huo.

**Keywords:** *Lactuca sativa*, TCP transcription factors, miR319, *LsTCP4*, plant regeneration

## Abstract

Plant regeneration is a major determinant of transformation and genome-editing efficiency, yet the endogenous regulatory networks controlling regenerative competence in horticultural crops remain incompletely understood. The miR319–TCP module regulates multiple developmental processes in plants, but its function in lettuce regeneration has not been defined. Here, we performed a genome-wide analysis of the TEOSINTE BRANCHED1/CYCLOIDEA/PROLIFERATING CELL FACTOR (TCP) gene family in lettuce (*Lactuca sativa*). Thirty-three *LsTCP* genes were identified and classified into Class I/PCF, Class II/CIN, and Class II/CYC/TB1 groups. Five CIN-class genes, *LsTCP2*, *LsTCP3*, *LsTCP4*, *LsTCP10*, and *LsTCP24*, were predicted as high-confidence miR319 targets and supported by degradome-based cleavage evidence.

*MIR319*-overexpression (OX319) explants showed enhanced de novo shoot regeneration, with 94.5% regeneration efficiency and 1.92 shoots per explant, whereas STTM-miR319 suppression (S319) explants showed reduced regeneration, with 28.5% regeneration efficiency and 0.36 shoots per explant. These phenotypes were associated with altered expression of several miR319-targeted CIN-TCP genes, particularly *LsTCP4*, *LsTCP10*, and *LsTCP24*. Disruption of *LsTCP4* increased regeneration efficiency to 91.4% and shoot production to 2.05 shoots per explant, resembling the regeneration-enhancing effect of miR319 overexpression. In contrast, disruption of the non-target CIN gene *LsTCP17* did not significantly affect regeneration under the tested conditions. Together, these results identify *LsTCP4* as a key miR319-responsive negative regulator of de novo shoot regeneration and highlight miR319-mediated repression of *LsTCP4* as a potential endogenous strategy for improving lettuce regeneration.

## Introduction

Efficient plant regeneration is a central requirement for genetic transformation, genome editing, and functional genomics in crops. *De novo* shoot regeneration requires differentiated cells to undergo extensive developmental reprogramming and establish new shoot meristematic identities in response to wound- and hormone-associated signals, particularly auxin and cytokinin (Bull and Michelmore, 2022; Ikeuchi *et al*., 2019). However, regeneration capacity varies widely among species, genotypes, explant types, and culture conditions, frequently limiting the recovery of transgenic or genome-edited plants. Identifying endogenous developmental regulators that influence regenerative competence is therefore important for improving plant biotechnology, particularly in horticultural crops with genotype-dependent tissue-culture responses (Jiang et al., 2026c).

Transcription factors are central regulators of plant growth and development because they coordinate gene-expression programs controlling cell proliferation, differentiation, organ formation, and environmental responses (Dhatterwal *et al*., 2024; Spitz and Furlong, 2012; Thilakarathne *et al*., 2025). Their activity is frequently refined through post-transcriptional regulation by microRNAs (miRNAs), which direct target-mRNA cleavage or translational repression, and thereby contribute to developmental timing and cell-fate control (Bartel, 2004; Rubio-Somoza and Weigel, 2011; Zhao *et al*., 2025). Among plant-specific transcription factor families, TEOSINTE BRANCHED1/CYCLOIDEA/PROLIFERATING CELL FACTOR (TCP) proteins are important regulators of growth and developmental patterning. TCP proteins contain a conserved non-canonical basic helix–loop–helix TCP domain that mediates DNA binding and protein interaction (Cubas *et al*., 1999; Viola *et al*., 2023). Based on sequence variation within this domain, TCP proteins are classified into Class I/PCF and Class II proteins, the latter of which comprises the CINCINNATA (CIN) and CYCLOIDEA/TEOSINTE BRANCHED1 (CYC/TB1) clades (Martín-Trillo and Cubas, 2010; Nicolas and Cubas, 2016; Yin *et al*., 2018). Through their effects on cell proliferation, differentiation, organ boundaries, and hormone-associated signaling, TCP proteins influence leaf morphology, shoot architecture, floral development, senescence, and other developmental processes (Cubas *et al*., 1999; Nicolas and Cubas, 2016; Viola *et al*., 2023).

The miR319–TCP module is one of the best-characterized regulatory systems controlling CIN-TCP activity. In *Arabidopsis thaliana*, miR319 directs cleavage of a subset of CIN-class TCP transcripts, including *TCP2*, *TCP3*, *TCP4*, *TCP10*, and *TCP24* (Palatnik *et al*., 2003). These miR319-regulated TCPs contribute to lateral-organ development by restricting excessive cell proliferation and influencing boundary-associated developmental programs, including *CUP-SHAPED COTYLEDON* gene expression (Koyama *et al*., 2017). Altered miR319–TCP activity affects leaf shape, leaf maturation, petal growth, shoot development, and jasmonate-associated processes in Arabidopsis (Bresso *et al*., 2018; Koyama *et al*., 2017; Nag *et al*., 2009; Schommer *et al*., 2008). The developmental importance of CIN-TCP regulation is conserved across species (Lan and Qin, 2020); however, its phenotypic effects vary across developmental contexts. For example, loss of the CIN-like gene *CINCINNATA* causes altered leaf curvature in *Antirrhinum majus* (Nath *et al*., 2003), whereas miR319-mediated regulation of the CIN-TCP *LANCEOLATE* is required for compound-leaf development in tomato (Ori *et al*., 2007). In rice, sweet potato, switchgrass, cotton, and *Medicago truncatula*, miR319-regulated TCP genes have also been associated with leaf architecture, photosynthetic performance, stress responses, fiber development, ethylene-related regulation, and nodulation (Cao *et al*., 2020; Liu *et al*., 2019; Ren *et al*., 2021; Wang *et al*., 2018; Yang *et al*., 2013). These findings indicate that the miR319–TCP module is evolutionarily conserved but can be deployed in species- and organ-specific developmental contexts.

Recent evidence also connects miR319-regulated TCP activity with plant regeneration. In *Arabidopsis*, reduced miR319 accumulation in the *hen1* background increased *TCP3* and *TCP4* expression and impaired *de novo* shoot regeneration (Yang *et al*., 2020). Genetic and molecular analyses further showed that *TCP4* can activate the cytokinin-response inhibitor *ARR16*, linking the miR319–TCP module to cytokinin responsiveness during shoot regeneration (Efroni *et al*., 2013; Yang *et al*., 2020). These findings provide a mechanistic precedent for the role of miR319-targeted CIN-TCPs in regenerative competence. However, whether this regulatory relationship is conserved in crop species remains largely unknown. In particular, the relative contributions of miR319-targeted and non-target CIN-TCP family members to regeneration have not been systematically examined.

Lettuce (*Lactuca sativa* L.) is an economically important leafy vegetable crop in which leaf morphology, size, developmental timing, and regenerative capacity directly influence production, breeding, and biotechnology applications (Akçakale Kaba *et al*., 2025; Chen *et al*., 2018; Huo *et al*., 2025; Lebeda *et al*., 2025). Although TCP proteins are likely to be important regulators of these traits, their functions in lettuce remain incompletely understood. Previous work identified *LsTCP4* as a candidate gene associated with the transition between Salinas- and Empire-type leaf morphology in crisphead lettuce (Seki *et al*., 2020). In addition, heterologous expression of *LsTCP13* and *LsTCP17* in *Arabidopsis* promoted flowering, suggesting that individual lettuce CIN-TCP genes retain developmental regulatory activity (Yun *et al*., 2023). These observations highlight the need for a systematic, genome-wide framework to define the lettuce TCP family, identify candidate miR319-regulated members, and prioritize genes for functional analysis. Genome-wide analyses in several crop species, including rice, cotton, *M. truncatula*, and sweet potato, have provided important frameworks for identifying TCP family members, predicting miR319 targets, and prioritizing genes for functional characterization (Ren *et al*., 2021; Wang *et al*., 2018; Yao *et al*., 2007; Yin *et al*., 2018). Similar genome-wide studies of other transcription-factor families in lettuce, including AP2/ERF, ARF, Aux/IAA, YABBY, and SPL families, have expanded understanding of developmental and stress-associated regulatory networks in this crop (Hu *et al*., 2022; Luo *et al*., 2023; Park *et al*., 2023; Zhang *et al*., 2025). Notably, our previous study showed that the miR156–SPL module regulates lettuce development and regeneration, supporting the broader importance of miRNA-mediated transcription-factor networks in lettuce regenerative competence (Jiang *et al*., 2026b).

Despite these advances, the TCP gene family has not been comprehensively characterized in lettuce, and the miR319–TCP regulatory module has not been functionally evaluated in lettuce regeneration. It is also unclear whether all CIN-TCP members contribute similarly to regeneration or whether miR319-targeted and non-target CIN-TCP genes have distinct functional roles. In this study, we performed a genome-wide identification and characterization of the lettuce TCP family, including phylogenetic classification, conserved motif and gene-structure analysis, chromosomal distribution, collinearity, synteny, promoter cis-element composition, and tissue-expression profiling. We further identified candidate miR319-targeted CIN-TCP genes using target prediction and degradome validation. To evaluate the functional significance of the miR319–TCP module in regeneration, we examined *MIR319* overexpression and STTM-miR319 suppression lines and compared the regeneration responses of the miR319-targeted *tcp4* mutant and the non-target CIN-class *tcp17* mutant. This study provides a genomic resource for the lettuce TCP family and identifies *LsTCP4* as a key miR319-responsive regulator of *de novo* shoot regeneration in lettuce.

## 2. Materials and methods

### 2.1. Plant materials and growth conditions

Lettuce (*Lactuca sativa* L.) cultivar ‘Salinas’ was used throughout this study. Wild-type (WT), *MIR319*-overexpression (OX319), STTM-miR319 (S319), *tcp4*, and *tcp17* plants were used for the indicated experiments. The *tcp4* and *tcp17* mutant lines used in this study were previously generated and sequence-validated in a companion study of miR319–TCP regulation of lettuce leaf senescence (Jiang et al., 2026a).

Seeds were surface sterilized in 70% (v/v) ethanol for 1 min, followed by 1% (v/v) sodium hypochlorite for 8–10 min, and rinsed thoroughly with sterile distilled water. Seeds were germinated on half-strength Murashige and Skoog (1/2 MS) medium supplemented with 1% (w/v) sucrose and 0.7% (w/v) agar, with the pH adjusted to 5.8 before autoclaving (Murashige and Skoog, 1962). Seedlings, regenerated plants, and established transgenic lines were maintained in a controlled-environment growth chamber at 22–24 °C under a 16-h light/8-h dark photoperiod with a photosynthetic photon flux density of approximately 100 μmol m⁻² s⁻¹. Unless otherwise stated, plants used for molecular and regeneration experiments were grown under identical conditions.

### 2.2. Genome-wide identification and characterization of *LsTCP* genes

The lettuce reference genome assembly and corresponding annotation files were retrieved from the NCBI Genome database and Phytozome (Reyes-Chin-Wo *et al*., 2017). Putative TCP proteins were identified using two complementary approaches. First, the Hidden Markov Model profile of the TCP domain (PF03634) was retrieved from the Pfam database and used to search the lettuce proteome with the HMMER module implemented in TBtools-II, using an E-value threshold of 1 × 10⁻⁵ (Chen *et al*., 2023). Second, the 24 *Arabidopsis* TCP protein sequences were used as queries for BLASTP searches against the lettuce proteome.

Redundant candidates and incomplete protein sequences were removed. The remaining candidates were validated for the presence of the conserved TCP domain using Pfam, InterProScan, SMART, and NCBI Conserved Domain Search (Jones *et al*., 2014). Verified genes were designated LsTCP1 to LsTCP33 based on phylogenetic relationships with Arabidopsis TCP homologs and retained nomenclature used for comparative analysis.

Protein length predicted molecular weight, theoretical isoelectric point, instability index, aliphatic index, and grand average of hydropathicity were calculated using ProtParam. Subcellular localization was predicted using CELLO (Yu *et al*., 2006). The resulting protein characteristics are listed in Supplementary Table 2.

### 2.3. Phylogenetic, structural, and evolutionary analyses of *LsTCP* genes

Full-length TCP protein sequences from *L. sativa*, *Arabidopsis thaliana*, *Oryza sativa*, *Solanum lycopersicum*, and *Helianthus annuus* were retrieved from the NCBI Genome and Phytozome databases. Multiple sequence alignment was conducted using ClustalW in MEGA X with default parameters. A neighbor-joining phylogenetic tree was constructed using the p-distance model, pairwise deletion of gaps, and 1,000 bootstrap replicates using MEGA X (Kumar *et al*., 2018). The tree was visualized using iTOL v6 (Letunic and Bork, 2024). LsTCP proteins were classified as Class I/PCF, Class II/CIN, or Class II/CYC/TB1 according to their clustering with functionally characterized Arabidopsis TCP proteins (Martín-Trillo and Cubas, 2010).

Conserved motifs were identified using MEME Suite v5.5.5 with the following parameters: maximum number of motifs, 10; motif width, 6–50 amino acids; and E-value threshold, 1 × 10⁻⁵ (Bailey *et al*., 2009). TCP-domain positions were annotated using Pfam and NCBI Conserved Domain Search. Exon–intron structures and untranslated regions were extracted from the lettuce genome annotation file. The phylogenetic tree, motif architecture, TCP-domain distribution, and gene structures were integrated and visualized using TBtools-II (Chen *et al*., 2023).

Physical positions of *LsTCP* genes were extracted from the lettuce genome annotation and mapped onto the nine lettuce chromosomes using TBtools-II (Chen *et al*., 2023). Intra-genomic collinearity analysis was performed using the MCScanX module implemented in TBtools-II (Chen *et al*., 2023; Wang *et al*., 2012). Collinear relationships among *LsTCP* loci were visualized using chromosome-scale and Circos plots (Krzywinski *et al*., 2009).

Inter-species synteny analysis was conducted between lettuce and *A. thaliana*, *H. annuus*, *O. sativa*, and *S. lycopersicum*. Reciprocal protein similarity searches and collinearity analyses were performed using MCScanX in TBtools-II (Chen *et al*., 2023; Wang *et al*., 2012). Syntenic blocks and TCP-related collinear gene pairs were visualized using the Multiple Synteny Plot and Advanced Circos modules of TBtools-II.

For homologous *LsTCP* pairs identified in collinearity analysis, synonymous substitution rates (Ks), nonsynonymous substitution rates (Ka), and Ka/Ks ratios were calculated using the Ka/Ks Calculator module in TBtools-II. Ka/Ks values below 1, equal to 1, and above 1 were interpreted as evidence consistent with purifying, neutral, and diversifying selection, respectively.

### 2.4. Promoter cis-element, predicted protein interaction, and Gene Ontology enrichment analyses

For each *LsTCP* gene, the 2,000-bp genomic region upstream of the predicted translation start codon was extracted from the lettuce genome and analyzed using PlantCARE (Lescot *et al*., 2002). Identified cis-regulatory elements were grouped into functional categories, including hormone-responsive, light-responsive, stress-responsive, meristem-related, wound-responsive, low-temperature-responsive, and cell-cycle-related elements. The number, category, and relative position of predicted cis-elements were visualized alongside the *LsTCP* phylogeny using TBtools-II (Chen *et al*., 2023).

Potential functional associations among LsTCP proteins were predicted using STRING with a medium-confidence interaction score threshold of 0.4 (Szklarczyk *et al*., 2019). The predicted interaction network was visualized using STRING. Proteins represented in the network were subjected to Gene Ontology enrichment analysis using ShinyGO (Xijin Ge *et al*., 2020). Biological-process terms with a false discovery rate (FDR) < 0.05 were retained for visualization. These analyses were used to identify candidate regulatory and functional associations among LsTCP proteins.

### 2.5. Identification, target prediction, and degradome validation of miR319

Mature miR319 sequences from *A. thaliana*, *L. sativa*, *S. lycopersicum*, and *O. sativa* were retrieved from miRBase and aligned using MEGA X (Kozomara *et al*., 2019; Kumar *et al*., 2018). Lettuce MIR319 loci were identified from the lettuce genome and designated *LsMIR319a* to *LsMIR319f*.

Potential miR319 target sites in *LsTCP* transcripts were predicted using psRNATarget and cross-validated using PmiREN (Dai *et al*., 2018; Guo *et al*., 2022). Genes predicted by both platforms were considered high-confidence candidate miR319 targets. Candidate target sites and their corresponding TCP-clade assignments are summarized in Table 1.

**Table 1.**
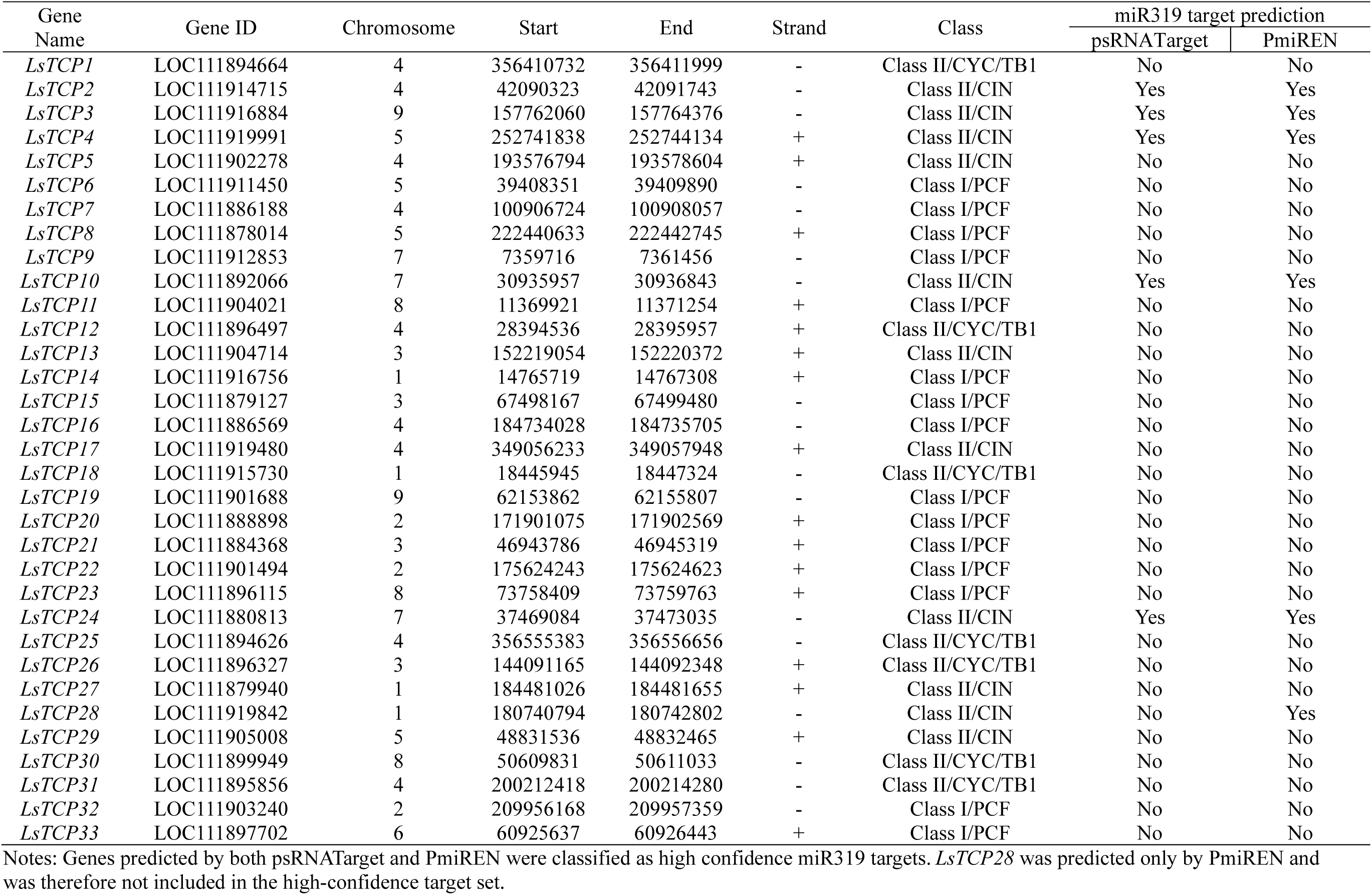
Genomic characteristics and miR319-target prediction of *LsTCP* genes in lettuce.

Publicly available lettuce degradome/PARE libraries were analyzed using CleaveLand4 to validate miR319-guided cleavage of predicted *LsTCP* targets (Addo-Quaye *et al*., 2009). Candidate cleavage signals were retained when degradome reads mapped to the canonical cleavage position between nucleotides 10 and 11 of the miR319 complementary site. Target plots were generated for *LsTCP2*, *LsTCP3*, *LsTCP4*, *LsTCP10*, and *LsTCP24*, and degradome categories were assigned according to CleaveLand4 classification criteria.

### 2.6. Tissue expression profiling and quantitative real-time PCR analysis

RNA-seq data from lettuce leaf, stem, and root tissues were obtained from the NCBI Gene Expression Omnibus under accession number GSE264560. Expression values for *LsTCP* genes were extracted from the published dataset. Genes with RPKM values < 2 across all sampled tissues were treated as lowly expressed. Expression values were normalized by gene for heatmap visualization, and hierarchical clustering was performed using TBtools-II (Chen *et al*., 2023).

For qRT-PCR analysis, cotyledon explants were collected after 2 weeks of culture on shoot-induction medium. Samples were immediately frozen in liquid nitrogen and stored at −80 °C until RNA extraction. Total RNA was isolated from approximately 100 mg tissue using RNAzol RT according to the manufacturer’s instructions. RNA concentration and purity were assessed using a NanoDrop spectrophotometer, and genomic DNA was removed using RNase-free DNase I.

First-strand cDNA was synthesized from 1 μg total RNA using the QuantiTect Reverse Transcription Kit in a 20-μL reaction volume. The resulting cDNA was diluted 20-fold before qRT-PCR. Mature miR319 was reverse transcribed using a stem-loop primer-based method. Gene-specific primers are listed in Supplementary Table 1, and *LsUBQ5* was used as the internal reference gene.

qRT-PCR was conducted using SYBR Green chemistry in 10-μL reactions containing 5 μL of 2× SYBR Green Master Mix, 0.5 μL each of forward and reverse primers (10 μM), 1 μL diluted cDNA template, and 3 μL nuclease-free water. The amplification program consisted of an initial denaturation step at 95 °C for 10 min, followed by 40 cycles of 95 °C for 15 s, 60 °C for 30 s, and 72 °C for 30 s. Melt-curve analysis was performed after amplification to verify primer specificity. Each biological sample was analyzed using three technical replicates, and three independent biological replicates were included per genotype. Relative transcript abundance was calculated using the 2⁻ΔΔCt method (Livak and Schmittgen, 2001).

### 2.7. Construction and Agrobacterium-mediated transformation of *MIR319*-related vectors

For *MIR319* overexpression, the lettuce *MIR319* precursor sequence was cloned into the binary vector pOX135 under the control of the 2× CaMV35S promoter. For miR319 suppression, a short tandem target mimic construct (STTM-miR319) was designed according to the strategy described by Tang *et al*. (2012). The STTM-miR319 cassette contained two imperfect miR319-binding sites separated by a 48-nt spacer, with a three-nucleotide bulge positioned around the predicted cleavage site to prevent target cleavage while preserving miRNA binding. The cassette was cloned into pOX135 under the control of the 2× CaMV35S promoter (Jiang et al., 2026a).

All constructs were confirmed by Sanger sequencing and introduced into *Agrobacterium tumefaciens* strain EHA105. The bacterial strain was grown overnight in LB medium containing spectinomycin (100 mg L⁻¹) and rifampicin (50 mg L⁻¹) at 28 °C with shaking at 180 rpm. A secondary culture was prepared in fresh antibiotic-containing LB medium and grown to an OD₆₀₀ of 0.3–0.5. Bacterial cells were collected by centrifugation and resuspended in liquid MS medium supplemented with 100 μM acetosyringone.

Cotyledons excised from 12- to 14-day-old seedlings were immersed in the Agrobacterium suspension for 10 min with gentle agitation. Inoculated explants were blotted dry on sterile filter paper and co-cultivated on MS medium containing 30 g L⁻¹ sucrose in the dark at 25 °C for 2 d. Explants were subsequently transferred to shoot-induction medium containing MS basal salts, 2 mg L⁻¹ 6-benzylaminopurine, 0.1 mg L⁻¹ naphthaleneacetic acid, 100 mg L⁻¹ kanamycin, and 100 mg L⁻¹ Timentin. Explants were subcultured onto fresh medium every 2 weeks.

Regenerated shoots were transferred to half-strength MS medium containing 100 mg L⁻¹ kanamycin for rooting. GFP fluorescence was used for preliminary identification of transgenic events, and transgene insertion was subsequently confirmed by PCR using construct-specific primers. Confirmed OX319 and S319 lines were propagated for molecular characterization and regeneration assays.

### 2.8. In vitro *de novo* shoot regeneration assay

Cotyledon explants of uniform size were collected from WT, S319, OX319, tcp4, and tcp17 plants. Explants were cultured on shoot-induction medium containing MS basal salts, 2 mg L⁻¹ 6-benzylaminopurine, and 0.1 mg L⁻¹ naphthaleneacetic acid under the growth conditions described above.

Regeneration was monitored weekly for 3 weeks, and representative whole-plate and close-up images were captured at weeks 1, 2, and 3. At the final scoring time point, regeneration capacity was evaluated using regeneration efficiency and shoots per explant. An explant was scored as regenerated when it produced at least one visible shoot ≥ 5 mm in length. Both metrics were calculated using the total number of cultured explants as the denominator.

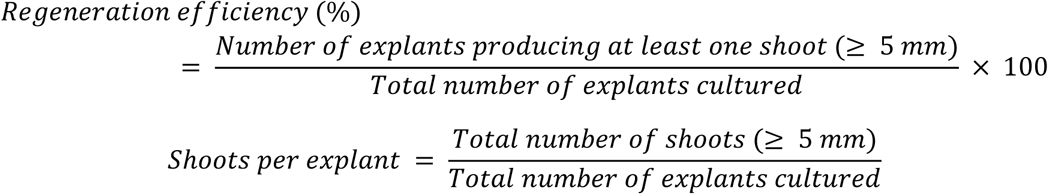

Each genotype was evaluated using 10 independent biological replicates. Each replicate consisted of a separate culture plate containing approximately 12–17 cotyledon explants, depending on explant availability and uniformity at the time of excision (Jiang et al., 2026c).

### 2.9. Statistical analysis

All experiments included at least three independent biological replicates. Regeneration assays included 10 biological replicates per genotype. Data are presented as means ± standard deviation unless otherwise stated in the figure legends.

Statistical analyses were conducted using GraphPad Prism 10. For comparisons among more than two genotypes, one-way analysis of variance, followed by Tukey’s honestly significant difference (HSD) post hoc test, was used. Pairwise comparisons were analyzed using two-tailed Student’s *T*-tests where appropriate. Statistical significance was defined as *P* < 0.05. Exact sample sizes, statistical tests, and significance annotations are provided in the corresponding figure legends.

## 3. Results

### 3.1. Genome-wide identification and classification of the lettuce TCP family

A genome-wide search combining HMMER screening using the TCP-domain profile and BLASTP searches with *Arabidopsis* TCP proteins identified 33 non-redundant TCP genes in the lettuce genome. After confirmation of the conserved TCP domain, the genes were designated *LsTCP1* to *LsTCP33* (Table 1). The encoded proteins ranged from 126 amino acids for LsTCP22 to 544 amino acids for LsTCP8, with predicted molecular weights ranging from 14.10 to 58.99 kDa and theoretical isoelectric points ranging from 5.25 to 9.90. All identified proteins contained the conserved TCP domain, whereas their additional motif composition, gene structure, and predicted physicochemical properties varied among family members (Table 1; Supplementary Table 2).

To classify the lettuce TCP family, a phylogenetic tree was constructed using full-length TCP protein sequences from lettuce, *Arabidopsis*, rice, tomato, and sunflower (Fig. 1A). The 33 LsTCP proteins were assigned to the three canonical TCP groups: Class I/PCF, Class II/CIN, and Class II/CYC/TB1. The Class I/PCF group was the largest, containing 15 members (*LsTCP6*, *LsTCP7*, *LsTCP8*, *LsTCP9*, *LsTCP11*, *LsTCP14*, *LsTCP15*, *LsTCP16*, *LsTCP19*, *LsTCP20*, *LsTCP21*, *LsTCP22*, *LsTCP23*, *LsTCP32*, and *LsTCP33*). The Class II/CIN clade contained 11 members (*LsTCP2*, *LsTCP3*, *LsTCP4*, *LsTCP5*, *LsTCP10*, *LsTCP13*, *LsTCP17*, *LsTCP24*, *LsTCP27*, *LsTCP28*, and *LsTCP29*), whereas the Class II/CYC/TB1 clade contained seven members (*LsTCP1*, *LsTCP12*, *LsTCP18*, *LsTCP25*, *LsTCP26*, *LsTCP30*, and *LsTCP31*) (Fig. 1A; Table 1).

**Figure 1.**
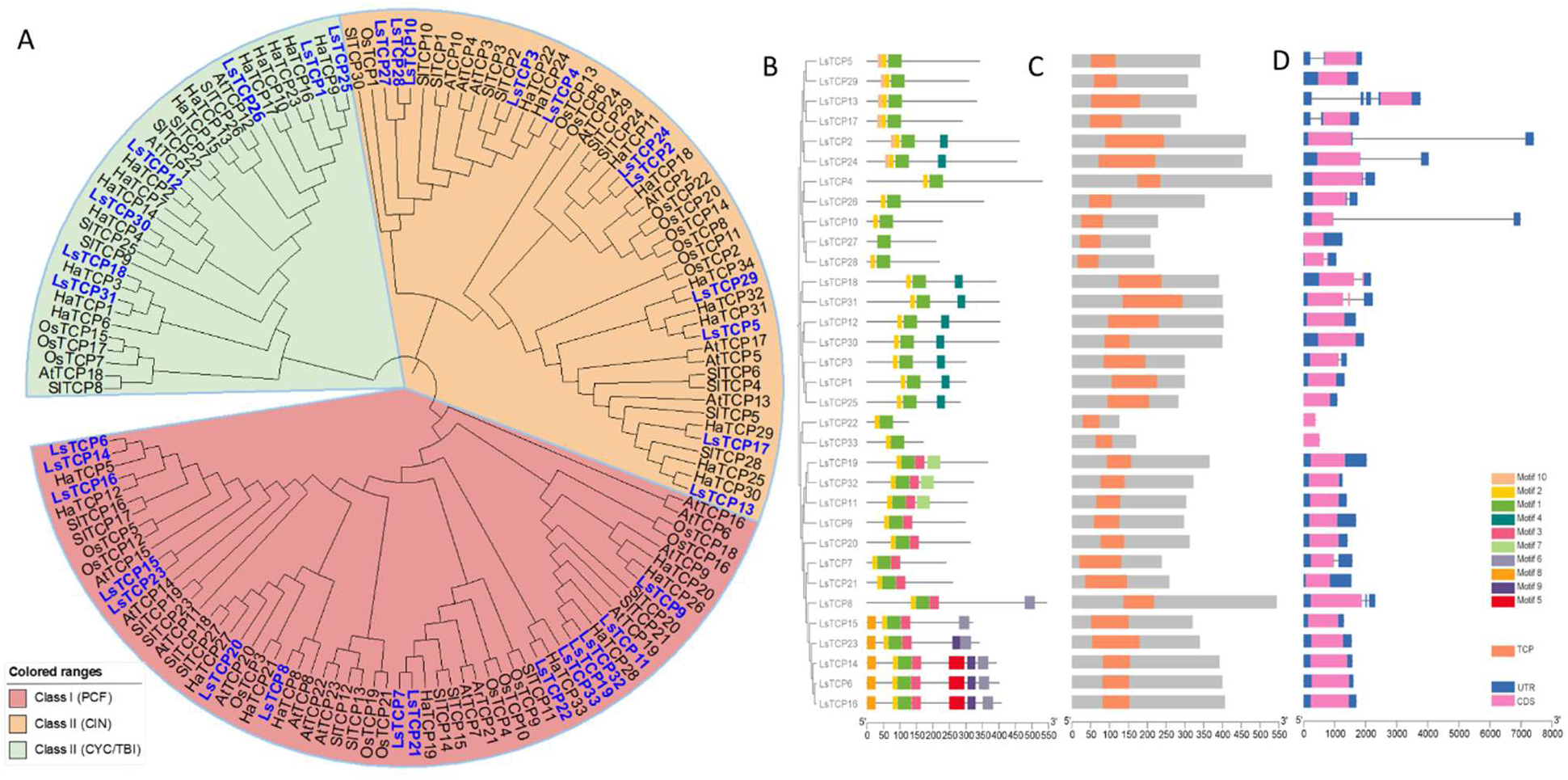
Phylogenetic classification, conserved motifs, TCP domains, and gene structures of TCP proteins. **A)** Phylogenetic tree of TCP proteins from *Lactuca sativa*, *Arabidopsis thaliana*, *Oryza sativa*, *Solanum lycopersicum*, and *Helianthus annuus*. Full-length amino acid sequences of 33 LsTCPs, 24 AtTCPs, 22 OsTCPs, 30 SlTCPs, and 34 HaTCPs were used to construct a neighbor-joining phylogenetic tree with 1,000 bootstrap replicates using MEGA X. TCP proteins were classified into three major groups: Class I/PCF, Class II/CIN, and Class II/CYC/TB1, which are highlighted with different background colors. LsTCP proteins are labeled in blue. **B–D)** Integrated analysis of LsTCP phylogeny, conserved motifs, conserved domains, and gene structures. B) Conserved motif composition of LsTCP proteins, with different colored boxes representing distinct motifs. C) Distribution of the conserved TCP domain within LsTCP proteins. D) Exon–intron organization of LsTCP genes. UTRs, coding sequences, and introns are represented by blue boxes, pink boxes, and black lines, respectively.

Five CIN-class genes, *LsTCP2*, *LsTCP3*, *LsTCP4*, *LsTCP10*, and *LsTCP24*, were predicted as high-confidence miR319 targets by both psRNATarget and PmiREN (Table 1). By contrast, *LsTCP13* and *LsTCP17*, although belonging to the CIN clade, lacked miR319 target predictions from both platforms. Thus, the lettuce CIN clade contains both candidate miR319-targeted and non-target TCP members.

### 3.2. Conserved structural features and evolutionary relationships of *LsTCP* genes

Conserved motif, domain, and gene-structure analyses were performed together with the phylogenetic classification to examine structural relationships within the LsTCP family (Fig. 1B–D). MEME analysis identified 10 conserved motifs among LsTCP proteins (Fig. 1B; Supplementary Fig. 1). Motif 1 was detected in all 33 proteins, whereas motif 2 was present in 32 proteins. The distribution of these conserved motifs was consistent with the TCP domain identified by Pfam and CD-Search analyses (Fig. 1C). Proteins grouped within the same phylogenetic clade generally displayed similar motif arrangements, whereas several motifs were restricted to specific clades, indicating structural differentiation among PCF, CIN, and CYC/TB1 members.

Gene-structure analysis showed variation in exon–intron organization across the LsTCP family (Fig. 1D). Most *LsTCP* genes contained few introns, with intron number ranging from zero to two. The majority of genes were intron poor, whereas *LsTCP4* and *LsTCP10* contained relatively long intronic regions. These data indicate that the LsTCP family is structurally conserved at the TCP-domain level but exhibits variation in motif composition and gene architecture among clades.

The 33 *LsTCP* genes were unevenly distributed across all nine lettuce chromosomes (Fig. 2A; Table 1). Chromosome 4 contained nine *LsTCP* genes, followed by chromosomes 1, 3, and 5, each containing four genes. Chromosomes 2, 7, and 8 each contained three genes, chromosome 9 contained two genes, and chromosome 6 contained one gene.

**Figure 2.**
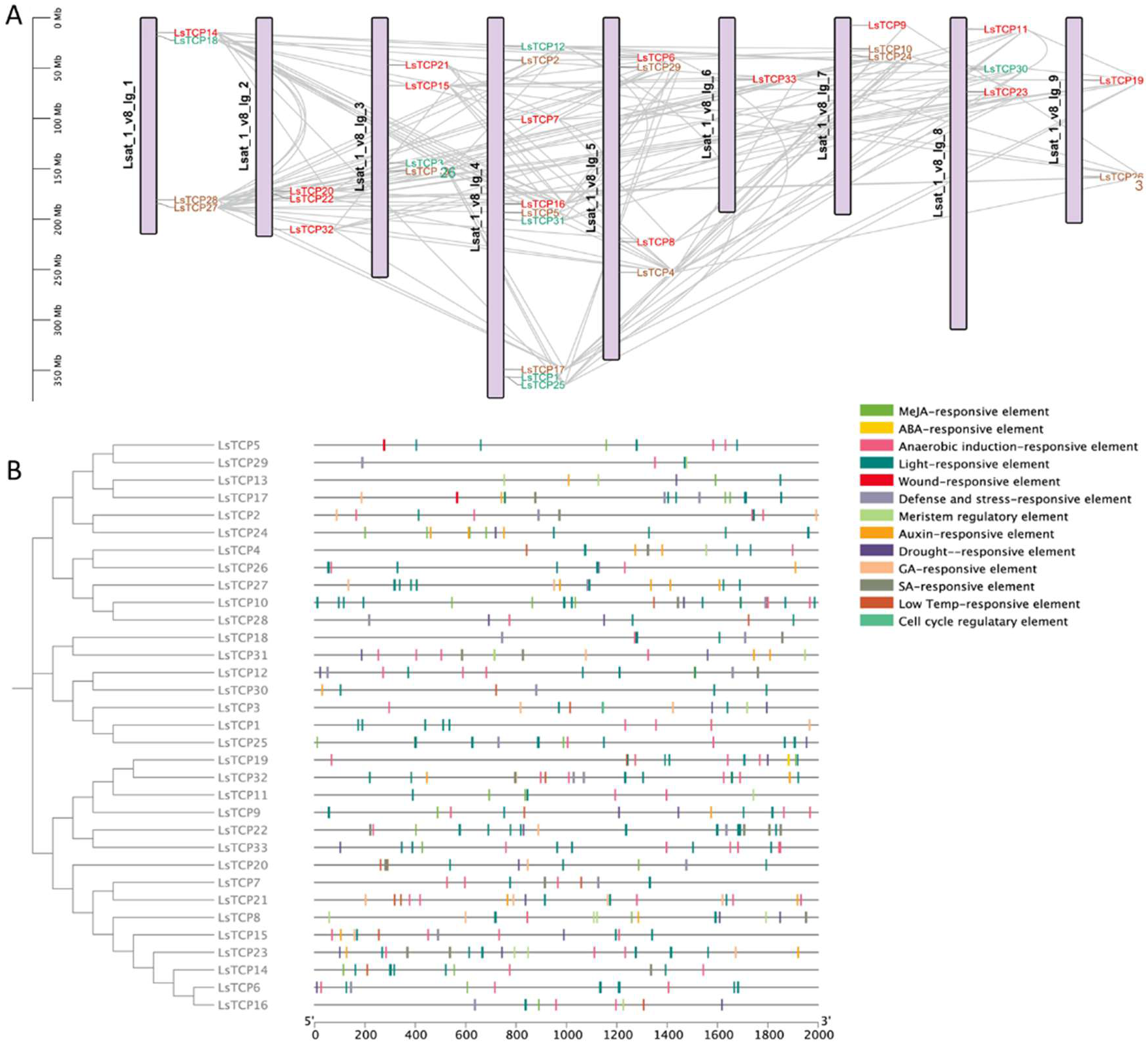
Chromosomal distribution, duplication relationships, and promoter cis-element analysis of *LsTCP* genes in lettuce. **A)** Chromosomal localization and collinearity analysis of the 33 *LsTCP* genes across nine lettuce chromosomes. The physical positions of *LsTCP* genes are shown according to chromosome length, with the scale indicated in megabases (Mb). Gray lines indicate putative segmental duplication or collinear relationships among *LsTCP* genes, suggesting potential evolutionary expansion of the TCP gene family in lettuce. Gene names are colored according to their phylogenetic classification or subgroup assignment. **B)** Cis-regulatory element analysis of the 2,000-bp upstream promoter regions of *LsTCP* genes. The left panel shows the phylogenetic relationships among LsTCP proteins, and the right panel shows the distribution of predicted cis-elements in each promoter region. Different colored vertical bars represent distinct regulatory elements, including hormone-responsive, light-responsive, stress-responsive, meristem-related, wound-responsive, low-temperature-responsive, and cell cycle regulatory elements. The position of each bar indicates its relative location within the promoter region from 5′ to 3′. The scale bar represents promoter length in base pairs.

Intra-genomic collinearity analysis identified multiple collinear relationships among *LsTCP* loci, indicating that duplication events contributed to the present organization of the family (Figs. 2A and 3A). Pairwise Ka/Ks analysis showed that most examined homologous *LsTCP* pairs had Ka/Ks ratios below 1, with values ranging from 0.058 to 1.067 (Supplementary Table 3). Only the *LsTCP26–LsTCP14* pair showed a Ka/Ks value slightly above 1.0. These results are consistent with predominant purifying selection acting on most examined *LsTCP* homologous pairs.

**Figure 3.**
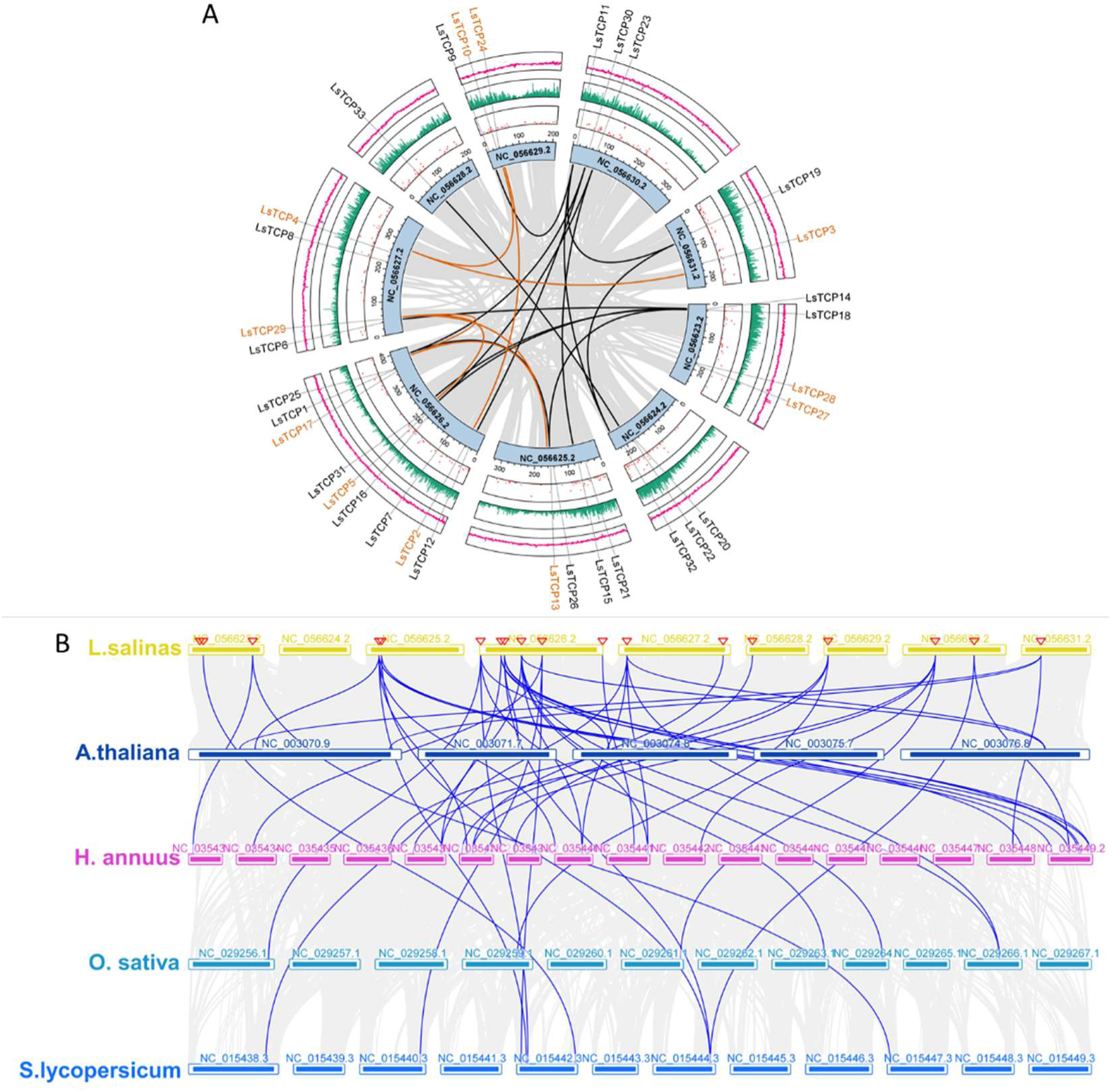
Intra- and inter-genomic collinearity analysis of lettuce *TCP* genes. **A)** Circos plot showing the chromosomal distribution, genomic features, and intra-genomic collinearity of *LsTCP* genes in the lettuce genome. The inner circle represents the nine lettuce chromosomes, with chromosome IDs indicated on each segment. Gray links indicate genome-wide collinear relationships, while colored links highlight putative duplicated or collinear *LsTCP* gene pairs. From the inner to outer tracks, the plot shows N-ratio distribution, GC content, gene density, and the physical positions of *LsTCP* genes. Gene names are shown around the outer circle, with selected *LsTCP* genes highlighted in orange. **B)** Inter-genomic synteny analysis of *TCP* genes between lettuce and four representative plant species, including *Arabidopsis thaliana*, *Helianthus annuus*, *Oryza sativa*, and *Solanum lycopersicum*. Chromosomes or genomic scaffolds from *L. sativa*, *A. thaliana*, *H. annuus*, *O. sativa*, and *S. lycopersicum* are shown as yellow, dark blue, pink, cyan, and blue boxes, respectively. Gray background lines indicate genome-wide syntenic blocks, whereas blue lines represent syntenic *TCP* gene pairs between lettuce and the other species, reflecting the evolutionary conservation and divergence of the *TCP* gene family.

Inter-species synteny analysis identified TCP-related collinear pairs between lettuce and Arabidopsis, sunflower, rice, and tomato (Fig. 3B). More extensive syntenic relationships were observed between lettuce and the eudicot species, particularly sunflower, whereas fewer collinear TCP loci were detected between lettuce and rice. This pattern was consistent with the closer evolutionary relationship between lettuce and sunflower, both of which belong to the Asteraceae family.

### 3.3. Predicted regulatory and functional features of the *LsTCP* family

To identify candidate regulatory features of the LsTCP family, the 2,000-bp promoter regions upstream of the translation start sites were analyzed using PlantCARE (Fig. 2B; Supplementary Table 4). A total of 516 predicted cis-regulatory elements were identified across the 33 *LsTCP* promoters. These elements included light-responsive, hormone-responsive, stress-responsive, meristem-related, wound-responsive, low-temperature-responsive, and cell-cycle-related motifs.

The most abundant individual predicted elements were associated with anaerobic induction, light responsiveness, methyl jasmonate responsiveness, and abscisic acid responsiveness. When grouped by function, light-responsive and hormone-responsive elements represented the most abundant categories. Auxin-responsive, meristem-related, and wound-responsive elements were also identified in multiple promoters. The composition and number of predicted cis-elements differed among *LsTCP* genes, providing candidate regulatory features for further functional investigation.

A predicted protein–protein interaction network was generated using STRING to examine potential associations among LsTCP proteins (Supplementary Fig. 2A). The resulting network contained multiple predicted associations among LsTCP family members, whereas some proteins showed relatively few predicted connections. Because these associations were generated computationally, they were treated as candidate functional relationships rather than direct evidence of protein interaction.

Gene Ontology enrichment analysis of proteins represented in the predicted network identified biological-process terms related to morphogenesis of branching structures, secondary shoot formation, inflorescence development, response to cytokinin, system development, shoot-system development, reproductive-structure development, and post-embryonic development (Supplementary Fig. 2B). These enriched terms were consistent with the established developmental functions of TCP transcription factors and provided hypotheses for subsequent functional analyses.

### 3.4. Public degradome data support miR319-guided cleavage of five CIN-class *LsTCP* transcripts in lettuce

To establish the miR319 target framework in lettuce, mature miR319 sequences from lettuce, *Arabidopsis*, tomato, and rice were aligned (Fig. 4A). The mature miR319 sequence was highly conserved among the examined species, particularly across the 5′ region, whereas limited sequence variation was present near the 3′ end. A genome-wide search identified six *MIR319* loci in lettuce, designated *LsMIR319a* to *LsMIR319f* (Fig. 4B).

**Figure 4.**
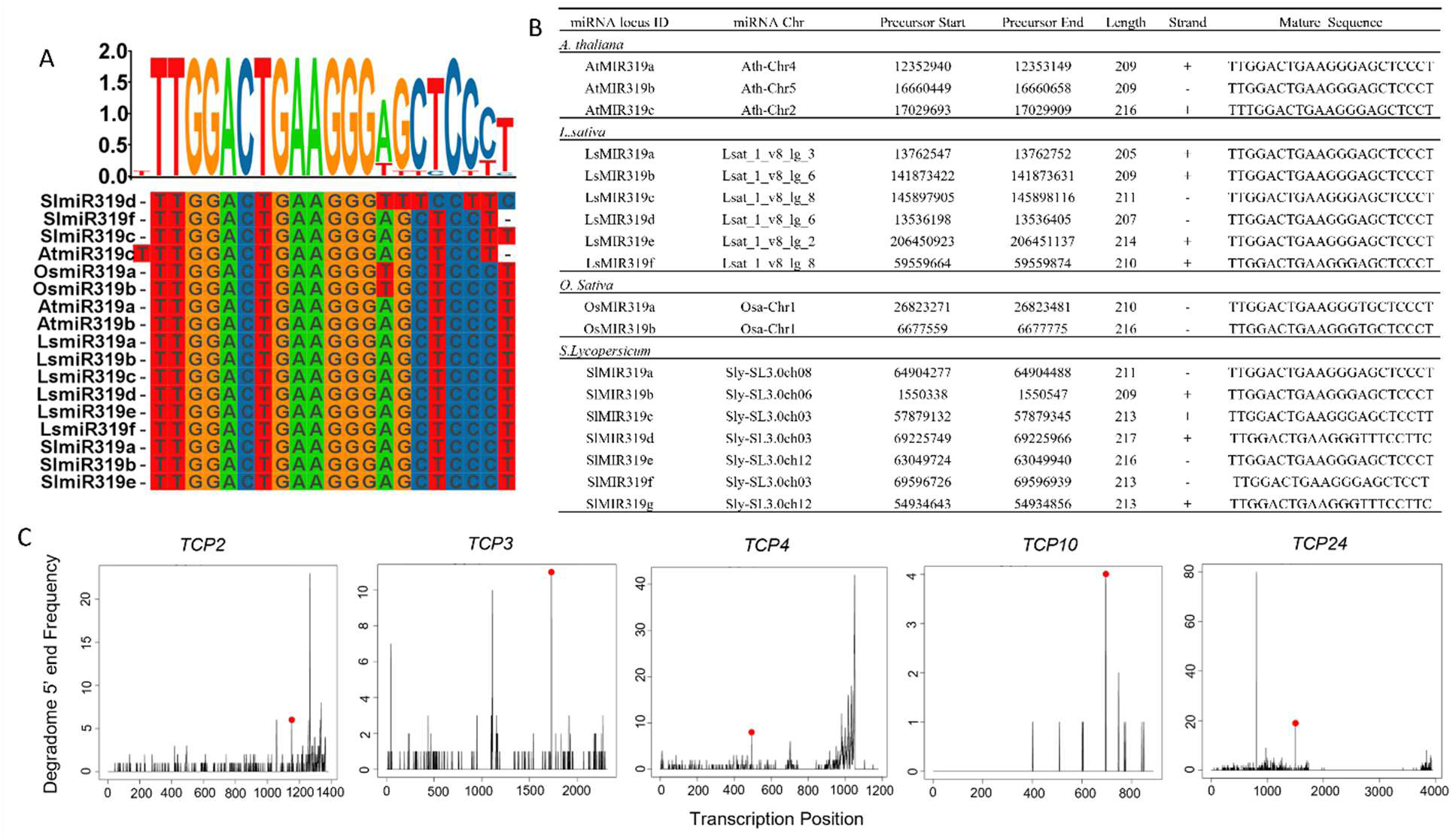
Conservation of miR319 and degradome-based validation of miR319-targeted *TCP* transcripts. **A)** Multiple sequence alignment of mature miR319 sequences from representative plant species, including *Arabidopsis thaliana*, *Lactuca sativa*, *Solanum lycopersicum*, and *Oryza sativa*. The sequence logo above the alignment shows nucleotide conservation, with letter height proportional to the frequency of each nucleotide at the corresponding position. **B)** Genomic locations and precursor-sequence information for MIR319 loci from *Arabidopsis thaliana*, *Lactuca sativa*, *Oryza sativa*, and *Solanum lycopersicum*. Conserved nucleotide positions are shown across species, supporting the evolutionary conservation of the miR319 family. **C)** Degradome analysis of predicted miR319-targeted *TCP* transcripts in lettuce. Each panel represents an individual *LsTCP* transcript. The x-axis indicates transcript position, and the y-axis indicates degradome read abundance. Red dots mark the predicted miR319-guided cleavage sites, providing degradome-supported evidence for cleavage of the indicated LsTCP transcripts.

Target prediction using both psRNATarget and PmiREN identified five high-confidence miR319 target candidates within the lettuce CIN clade: *LsTCP2, LsTCP3, LsTCP4, LsTCP10,* and *LsTCP24* (Table 1). To assess support for these predicted interactions, publicly available degradome data were analyzed for miR319-guided cleavage signals. Cleavage signals mapping to the expected miR319 complementary site were detected for all five transcripts (Fig. 4C). The predicted cleavage site coincided with the dominant degradome peak for *LsTCP3* and *LsTCP10*, which were classified as category 0 targets. In contrast, *LsTCP2, LsTCP4,* and *LsTCP24* showed category-2 cleavage signals at the predicted miR319 cleavage position. Together, these results support the identification of five directly cleaved CIN-class *LsTCP* transcripts as miR319 targets in lettuce.

Having established the target and non-target CIN-TCP framework, we next examined the tissue expression patterns of these genes and their transcriptional responses to altered miR319 activity.

### 3.5. Tissue expression and altered miR319 dosage reveal differential regulation of CIN-TCP genes

RNA-seq data from leaf, stem, and root tissues showed that the 33 *LsTCP* genes displayed distinct expression patterns across the three organs (Fig. 5A). Several genes, including *LsTCP20* and *LsTCP21*, showed relatively high expression across leaf, stem, and root tissues, whereas *LsTCP25, LsTCP26, LsTCP28,* and *LsTCP29* were expressed at lower levels under the analyzed conditions.

**Figure 5.**
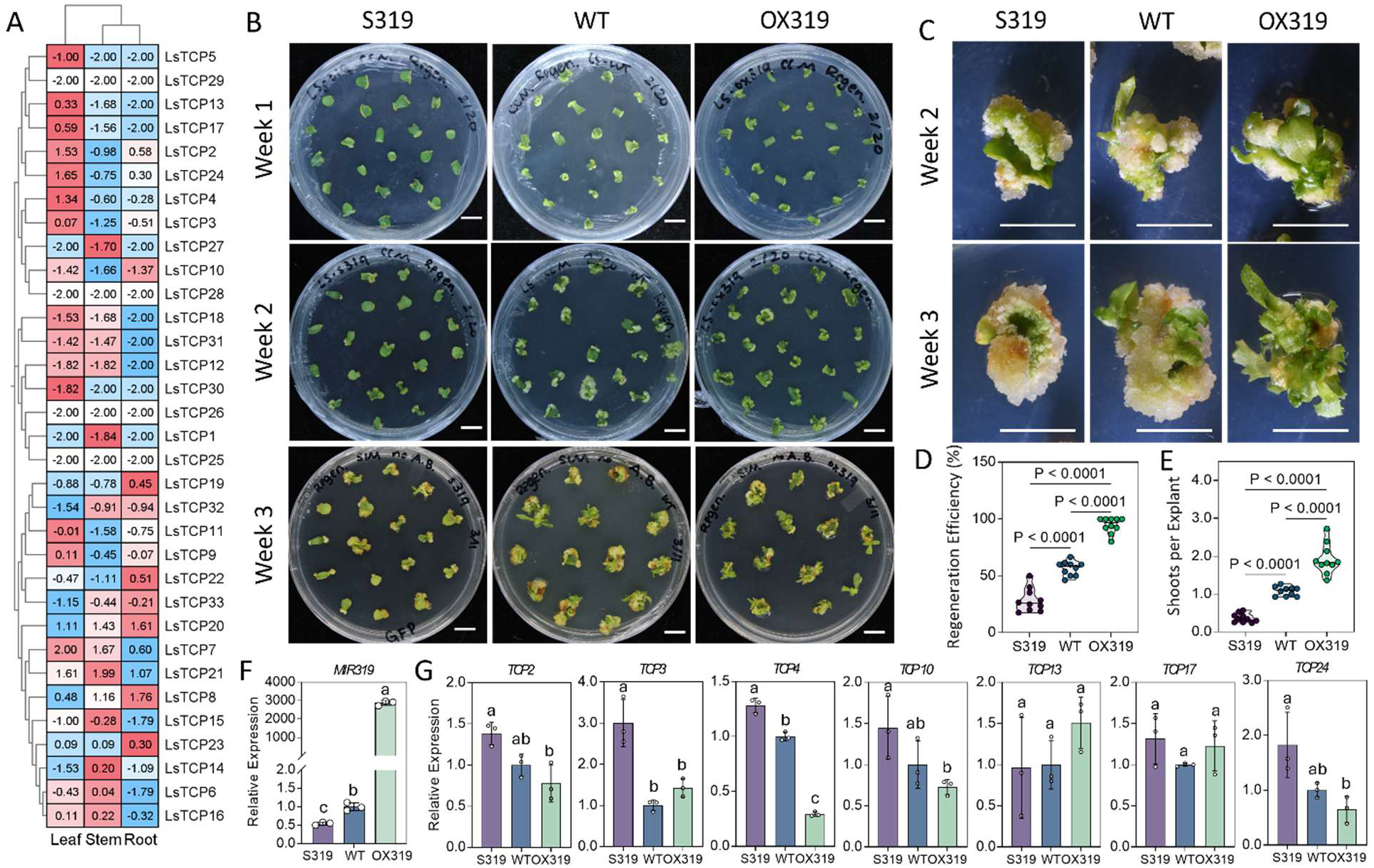
Tissue expression profiles and regeneration phenotypes associated with altered miR319 activity in lettuce. **A)** Tissue-specific expression profiles of *LsTCP* genes in leaf, stem, and root tissues. Expression values were obtained from NCBI GEO DataSets under accession number GSE264560 and are shown as a heatmap after normalization. Red and blue indicate relatively high and low expression levels, respectively. **B)** Representative regeneration phenotypes of wild-type, *STTM-miR319*, and *MIR319*-overexpression explants during *in vitro* culture at weeks 1, 2, and 3. **C)** Representative close-up images of regenerating explants at weeks 2 and 3, showing differences in callus formation and shoot regeneration among genotypes. **D, E)** Quantification of regeneration performance, including regeneration efficiency **(D)** and shoots per explant **(E)**. Each point represents one independent culture plate as a biological replicate; n = 10 biological replicates per genotype. Statistical significance was determined by one-way ANOVA followed by Tukey’s multiple-comparison test; pairwise adjusted *P* values are shown above the plots. **F, G)** Relative expression levels of miR319 **(F)** and selected *LsTCP* genes **(G)**, as determined by qRT-PCR. Data are presented as mean ± SD. Three independent biological replicates were analyzed, each with three technical replicates. Different letters indicate significant differences among genotypes according to one-way ANOVA followed by Tukey’s HSD test (*P* < 0.05). Scale bars: 5 mm.

Among the five directly cleaved miR319-targeted CIN genes, *LsTCP2, LsTCP4,* and *LsTCP24* showed preferential expression in leaf tissue. *LsTCP10* showed low expression in all three tissues, whereas *LsTCP3* showed relatively similar expression across leaf, stem, and root tissues. The non-target CIN genes *LsTCP13* and *LsTCP17* also showed detectable expression in leaf tissue, providing a basis for comparing miR319-targeted and non-target CIN members in subsequent experiments.

To determine how altered miR319 activity affects the expression of selected CIN-TCP genes, 2-week explants of STTM-miR319 (S319), *MIR319*-overexpression plants (OX319), and wild-type (WT) were examined by qRT-PCR (Fig. 5F, G). Mature miR319 abundance was approximately 2,875-fold higher in OX319 and 45.2% lower in S319 than in WT. Several validated miR319 targets showed expression changes consistent with altered miR319 dosage. In OX319, *LsTCP4, LsTCP10,* and *LsTCP24* were reduced to 29.4%, 72.7%, and 64.8% of WT expression, respectively; in S319, they increased to 1.28-, 1.45-, and 1.83-fold of WT levels, respectively.

*LsTCP2* was moderately reduced in OX319 and elevated in S319. *LsTCP3* showed a strong increase in S319 but did not show proportional repression in OX319. In contrast, the non-target CIN genes *LsTCP13* and *LsTCP17* did not show a consistent reciprocal response to altered miR319 activity. Overall, altered miR319 activity was associated with reciprocal expression changes in several directly cleaved CIN-class *LsTCP* genes, particularly *LsTCP4, LsTCP10,* and *LsTCP24*, whereas *LsTCP13* and *LsTCP17* showed no consistent reciprocal response. These patterns identified *LsTCP4* as a strongly miR319-responsive candidate for subsequent functional analysis and provided a non-target CIN-TCP comparison represented by *LsTCP17*.

### 3.6. miR319 dosage modulates de novo shoot regeneration, and LsTCP4 functions as a negative regulator

Having established the miR319 target framework and identified differential transcriptional responses among selected CIN-TCP genes, we next examined whether altered miR319 activity affects de novo shoot regeneration. Cotyledon explants from S319, WT, and OX319 plants were cultured on shoot-induction medium and monitored for three weeks (Fig. 5B, C). During the first week, explants from all three genotypes initiated callus at the cut edges. Differences in shoot formation became apparent from the second week onward. OX319 explants developed abundant green shoots and leafy structures by week three, whereas S319 explants showed limited shoot outgrowth. WT explants displayed an intermediate response.

Regeneration was quantified as the proportion of explants forming shoots and the number of shoots per explant. OX319 showed a mean regeneration efficiency of 94.5%, representing a 67.2% increase relative to WT, which showed a mean regeneration efficiency of 56.5% (Fig. 5D). In contrast, S319 showed a mean regeneration efficiency of 28.5%, representing a 49.5% reduction relative to WT. OX319 produced 1.92 shoots per explant, which was 78.9% higher than the WT value of 1.07 shoots per explant (Fig. 5E). S319 produced 0.36 shoots per explant, corresponding to a 66.1% reduction relative to WT. Together, these reciprocal phenotypes indicate that miR319 dosage positively influences de novo shoot regeneration in lettuce.

To determine whether individual CIN-TCP genes contribute to this regeneration response, we evaluated previously generated and sequence-validated CRISPR/Cas9-edited *tcp4* and *tcp17* lines (Jiang et al., 2026a). *LsTCP4* was selected because it is directly cleaved by miR319 and showed strong repression in OX319, whereas *LsTCP17* was selected as a CIN-clade gene lacking canonical miR319 target signatures and showing no consistent reciprocal response to altered miR319 activity. Representative regeneration responses are shown in Fig. 6A, B. During regeneration, *tcp4* explants developed multiple organized leafy shoots, whereas *tcp17* explants displayed an overall regeneration response similar to that of WT. The *tcp4* line showed a mean regeneration efficiency of 91.4%, corresponding to a 61.7% increase relative to WT, and produced 2.05 shoots per explant, representing a 91.1% increase relative to WT (Fig. 6C, D). In contrast, *tcp17* showed a mean regeneration efficiency of 55.2% and produced 0.98 shoots per explant; neither parameter differed significantly from WT under the tested regeneration conditions. Thus, disruption of *LsTCP4*, but not *LsTCP17*, enhanced regeneration under the tested conditions. The *tcp4* phenotype was similar to that of OX319, supporting *LsTCP4* as a miR319-responsive negative regulator of de novo shoot regeneration in lettuce.

**Figure 6.**
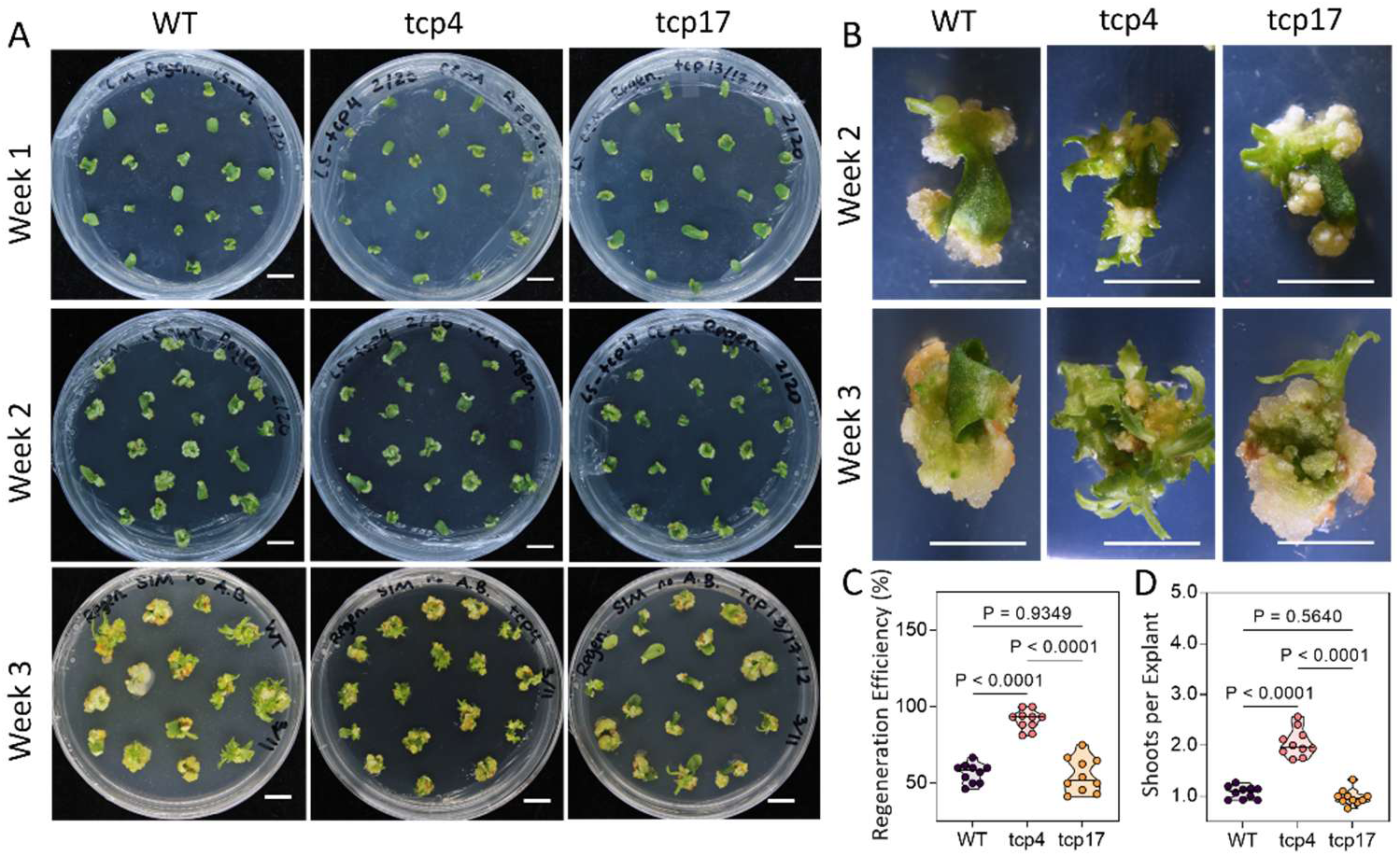
Loss of *LsTCP4*, but not *LsTCP17*, enhances de novo shoot regeneration in lettuce. **A)** Representative regeneration phenotypes of wild-type, *tcp4* edited, and *tcp17* edited lettuce explants during in vitro culture at weeks 1, 2, and 3. Whole-plate images show the progression of callus formation and shoot regeneration among the indicated genotypes. **B)** Representative close-up images of regenerating explants at weeks 2 and 3, showing genotype-dependent differences in callus development and shoot formation. **C, D)** Quantitative analysis of regeneration performance, including regeneration efficiency **(C)** and shoots per explant **(D)**. Each point represents one independent culture plate as a biological replicate; n = 10 biological replicates per genotype. Statistical significance was determined by one-way ANOVA followed by Tukey’s multiple-comparison test; pairwise adjusted P values are shown above the plots. Scale bars: 5 mm.

## 4. Discussion

### 4.1. Genome-wide characterization identifies a conserved miR319-responsive CIN-TCP module in lettuce

This study provides a genome-wide framework for the TCP gene family in lettuce and identifies a conserved miR319-responsive CIN-TCP module associated with de novo shoot regeneration. The 33 *LsTCP* genes were assigned to the Class I/PCF, Class II/CIN, and Class II/CYC/TB1 groups, consistent with the canonical organization of TCP families in other angiosperms. The conserved TCP domain was retained across all family members, whereas motif composition, intron–exon organization, chromosomal distribution, and tissue-expression profiles differed among genes and clades. Intra-genomic collinearity and Ka/Ks analyses further indicated that most examined homologous *LsTCP* gene pairs have been maintained under predominant purifying selection. Together, these data provide a foundation for distinguishing conserved family features from potentially divergent gene functions in lettuce.

A major outcome of the present study was the identification of five directly cleaved miR319-responsive CIN-class genes: *LsTCP2*, *LsTCP3*, *LsTCP4*, *LsTCP10*, and *LsTCP24*. These genes were predicted by both psRNATarget and PmiREN, and degradome analysis detected miR319-guided cleavage at the expected target site in all five transcripts. This result is consistent with the conserved regulation of a subset of CIN-TCP genes by miR319 in *Arabidopsis* and other plant species (Fang *et al*., 2021; Palatnik *et al*., 2003). Among these targets, *LsTCP2*, *LsTCP4*, and *LsTCP24* showed preferential expression in leaf tissue, whereas *LsTCP13* and *LsTCP17* lacked canonical miR319 target signatures and did not exhibit a consistent reciprocal expression response to altered miR319 activity. Thus, the lettuce CIN clade includes both directly miR319-regulated and non-target TCP members, providing a useful framework for examining their distinct developmental functions.

The promoter and predicted interaction analyses further suggested that *LsTCP* genes may participate in hormone- and development-associated regulatory networks. The enrichment of auxin-, methyl jasmonate-, abscisic acid-, meristem-, and wound-associated promoter motifs is compatible with the known involvement of TCP factors in developmental and hormone-responsive processes. Similarly, predicted Gene Ontology terms related to cytokinin response, shoot-system development, and secondary shoot formation are consistent with potential functions in regeneration-associated pathways. However, these analyses are computational predictions and should be considered hypothesis-generating rather than direct functional evidence. Wounding, cellular reprogramming, and hormone-dependent acquisition of shoot identity are central components of de novo shoot regeneration (Ikeuchi *et al*., 2019; Iwase *et al*., 2011), and direct molecular experiments will be required to determine how individual *LsTCP* genes participate in these processes.

### 4.2. miR319 promotes de novo shoot regeneration through repression of selected CIN-TCP genes, particularly *LsTCP4*

The functional experiments demonstrate that miR319 positively regulates de novo shoot regeneration in lettuce. Elevating miR319 activity in OX319 plants significantly increased both regeneration efficiency and shoot number per explant, whereas suppression of endogenous miR319 activity in S319 plants produced the opposite phenotype. These reciprocal regeneration responses were accompanied by altered expression of several validated miR319-targeted CIN-TCP genes, particularly *LsTCP4*, *LsTCP10*, and *LsTCP24*. Therefore, the regeneration phenotype associated with altered miR319 activity is consistent with changes in the expression of multiple CIN-TCP targets.

Among the tested targets, *LsTCP4* showed the strongest functional support as a negative regulator of regeneration. *LsTCP4* was strongly repressed in OX319 plants, and the *tcp4* mutant exhibited increased regeneration efficiency and shoot production relative to WT. The enhancement of regeneration in both OX319 and *tcp4* supports the conclusion that miR319 promotes regeneration, at least partly, through repression of *LsTCP4*. This interpretation is consistent with the established developmental roles of miR319-regulated TCP proteins in lateral-organ growth and leaf differentiation (Bresso *et al*., 2018; Koyama *et al*., 2017; Palatnik *et al*., 2003). Our findings are also consistent with evidence from *Arabidopsis*, in which miR319-mediated regulation of TCP3 and TCP4 contributes to de novo shoot regeneration. In the *Arabidopsis hen1* mutant, reduced miR319 abundance leads to elevated *TCP3* and *TCP4* expression, reduced cytokinin responsiveness, and impaired shoot regeneration. TCP4 directly activates *ARR16*, a type-A response regulator that negatively modulates cytokinin signaling during shoot regeneration (Yang *et al*., 2020). Earlier work also showed that CIN-TCP activity can reduce cytokinin sensitivity during leaf maturation through activation of type-A response regulators, including *ARR16* (Efroni *et al*., 2013). Together, these studies provide a plausible mechanistic framework for the lettuce regeneration phenotype observed here.

Nevertheless, the present study did not directly assess cytokinin responsiveness, *LsARR16* expression, shoot-meristem regulators, or the temporal transition from callus formation to shoot initiation. Therefore, it remains premature to conclude that the miR319–*LsTCP4* module promotes regeneration in lettuce through the same TCP–ARR16 cytokinin pathway described in *Arabidopsis*. Instead, our results establish *LsTCP4* as a major miR319-responsive negative regulator of lettuce regeneration and identify cytokinin signaling as a testable downstream hypothesis.

Because OX319 altered the expression of multiple miR319-targeted CIN-TCP genes, whereas the *tcp4* mutant disrupted only one target, the OX319 phenotype cannot be attributed exclusively to *LsTCP4*. The validated targets *LsTCP2*, *LsTCP3*, *LsTCP10*, and *LsTCP24,* therefore, remain important candidates for further functional characterization. Their effects may be partially redundant, additive, or dependent on the developmental stage of the explant, as has been observed for miR319-regulated TCP genes during leaf development in *Arabidopsis* (Bresso *et al*., 2018; Koyama *et al*., 2017). Combinatorial editing, multiplex CRISPR/Cas systems, and temporally controlled expression strategies will help determine whether simultaneous modulation of selected targets can further improve regeneration.

The present findings also complement our previous demonstration that the miR156–SPL module regulates regenerative competence in lettuce (Jiang et al., 2026b). Together, these studies indicate that lettuce regeneration is regulated by multiple endogenous microRNA–transcription factor modules rather than by a single developmental pathway. Determining whether miR156–SPL and miR319–TCP regulation converge on common cytokinin-responsive genes, meristem regulators, or wound-induced reprogramming pathways will be important for understanding how regenerative competence is established in lettuce explants.

### 4.3. Functional differentiation among CIN-TCP genes and implications for crop biotechnology

The comparison between *tcp4* and *tcp17* indicates that CIN-clade membership alone does not predict regeneration function. Unlike *tcp4*, *tcp17* did not differ significantly from WT in regeneration efficiency or shoots per explant under the tested conditions. This contrast is particularly informative because *LsTCP17* belongs to the CIN clade but lacks a canonical miR319 target signature and did not show a consistent reciprocal expression response to altered miR319 activity. Therefore, the enhanced regeneration observed in OX319 plants cannot be interpreted as a general consequence of reducing the activity of all CIN-TCP genes. Rather, the present data support functional differentiation among CIN-TCP members in the cotyledon-based lettuce regeneration system.

The absence of a detectable *tcp17* phenotype in this assay does not establish that *LsTCP17* lacks developmental function. Previous work showed that heterologous expression of *LsTCP13* and *LsTCP17* in *Arabidopsis* accelerated flowering, demonstrating that these lettuce CIN-TCP proteins retain developmental regulatory activity (Yun *et al*., 2023). In addition, *Arabidopsis* TCP family members can exhibit partially overlapping, tissue-specific, and condition-dependent functions (Bresso *et al*., 2018; Koyama *et al*., 2017). Thus, *LsTCP17* may function in developmental processes not captured by the final regeneration indices measured here, including responses that depend on explant identity, culture conditions, or developmental stage.

From a biotechnology perspective, the present data identify the miR319–*LsTCP4* branch, rather than CIN-TCP genes broadly, as a promising endogenous target for improving shoot regeneration during transformation and genome editing. Regeneration remains a major bottleneck in crop biotechnology, and lettuce regeneration is strongly influenced by genotype, explant source, and culture conditions (Altpeter *et al*., 2016; Bull and Michelmore, 2022). The enhanced regeneration observed in OX319 and *tcp4* lines indicates that reducing miR319-responsive *LsTCP4* activity can improve shoot recovery. However, stable manipulation of miR319-regulated TCP activity may entail developmental trade-offs because these factors have broad roles in leaf differentiation, organ growth, and developmental timing (Bresso *et al*., 2018; Fang *et al*., 2021; Palatnik *et al*., 2003).

Accordingly, practical application of this pathway may require transient, inducible, or regeneration-stage-specific manipulation. Potential approaches include transient miR319 delivery during tissue culture, inducible miR319 expression, temporary suppression of *LsTCP4*, or modification of regeneration-relevant regulatory regions rather than constitutive disruption of *LsTCP4*. These approaches remain to be tested, but they are consistent with broader efforts to use endogenous developmental regulators to improve crop transformation and genome editing while minimizing persistent developmental penalties (Altpeter *et al*., 2016).

### 4.4. Limitations and future perspectives

Several limitations should be considered when interpreting this study. First, the functional experiments were conducted in the ‘Salinas’ background using cotyledon explants and a single shoot-induction medium. The contribution of miR319 and individual TCP genes may vary among lettuce genotypes, explant types, developmental stages, and hormone combinations. Second, although the *tcp4* phenotype strongly supports a role for *LsTCP4*, the respective functions of the other directly cleaved miR319 targets remain unresolved. Third, the proposed cytokinin-related mechanism is inferred from studie*s* in *Arabidopsis* and has not been directly tested in lettuce. Finally, promoter motif, protein interaction, and Gene Ontology analyses provide useful hypotheses but do not establish regulatory interactions or molecular mechanisms.

Future work should therefore include phenotypic analysis of additional independent edited alleles, time-resolved transcript profiling during callus induction and shoot formation, cytokinin-response assays, and examination of lettuce homologs of *ARR16*, *WUSCHEL*, and other shoot-meristem regulators. Testing individual and multiplex edits of *LsTCP2*, *LsTCP3*, *LsTCP10*, and *LsTCP24* will help define target-specific and combinatorial contributions to regeneration. Evaluation across diverse lettuce genotypes and tissue-culture systems will also be essential to determine whether the miR319–TCP module can be broadly deployed to improve transformation and genome-editing efficiency.

## 5. Conclusion

This study establishes a genome-wide resource for the lettuce TCP family and identifies five directly cleaved miR319-responsive CIN-class genes: *LsTCP2*, *LsTCP3*, *LsTCP4*, *LsTCP10*, and *LsTCP24*. Functional analyses showed that elevated miR319 activity enhanced de novo shoot regeneration, whereas miR319 suppression reduced regeneration. Disruption of *LsTCP4* increased regeneration efficiency and shoot production and showed a regeneration-enhancing effect consistent with that of miR319 overexpression, identifying *LsTCP4* as a key miR319-responsive negative regulator of regeneration in lettuce. By contrast, disruption of the non-target CIN gene *LsTCP17* did not significantly alter regeneration under the tested conditions. Together, these results show that selected miR319-targeted CIN-TCP genes contribute to lettuce regenerative competence and provide a foundation for developing temporally controlled endogenous-regulator strategies to improve regeneration and transformation in lettuce.

## Author Contributions

T.J. and H.H. conceived and designed the study. T.J. performed the genome-wide identification and characterization of the lettuce TCP gene family, miR319 target prediction, degradome analysis, phylogenetic analysis, promoter cis-element analysis, collinearity/synteny analysis, data visualization, and manuscript preparation. S.E.T. contributed to plant tissue culture, regeneration assays, and molecular analyses. A.K. contributed to manuscript preparation. F.L. contributed to plant material preparation, mutant-line maintenance, and experimental support. T.J. analyzed the data and prepared the figures and tables. T.J. wrote the manuscript with input from all authors. H.H. supervised the project, provided funding and resources, and revised the manuscript. All authors read and approved the final manuscript.

## Data Availability Statement

Public RNA-seq data used in this study are available from the NCBI Gene Expression Omnibus under accession number GSE264560. All other data supporting the findings of this study are included in the article and its supplementary materials or are available from the corresponding author upon reasonable request.

## Acknowledgments

We thank Jaideep Chandranshu Cherukula for plant and facility management, and Keila Emily Rodriguez for assistance with previously generated lettuce genetic materials used in this project.

## Abbreviations

ARR16: Arabidopsis response regulator 16
BAP: 6-benzylaminopurine
CIN: CINCINNATA
CYC/TB1: CYCLOIDEA/TEOSINTE BRANCHED1
FDR: false discovery rate
GEO: Gene Expression Omnibus
GO: Gene Ontology
GRAVY: grand average of hydropathicity
HMM: hidden Markov model
HSD: honestly significant difference
Ka: nonsynonymous substitution rate
Ks: synonymous substitution rate
miRNA: microRNA
MS: Murashige and Skoog
NAA: naphthaleneacetic acid
OX319: MIR319-overexpression line
PARE: parallel analysis of RNA ends
PCF: PROLIFERATING CELL FACTOR
qRT-PCR: quantitative real-time PCR
S319: STTM-miR319 suppression line
STTM: short tandem target mimic
TCP: TEOSINTE BRANCHED1/CYCLOIDEA/PROLIFERATING CELL FACTOR
WT: wild type.

## Supplementary Materials

Supplementary Fig. 1. Conserved motif sequence logos of lettuce LsTCP proteins.

Supplementary Fig. 2. Predicted protein-association network and Gene Ontology enrichment analysis of lettuce LsTCP proteins.

Supplementary Table 1. Primers used for qRT-PCR analysis of selected *LsTCP* genes.

Supplementary Table 2. Physicochemical characterization of lettuce LsTCP proteins.

Supplementary Table 3. Ka/Ks analysis of selected *LsTCP* gene pairs.

Supplementary Table 4. Predicted cis-acting regulatory elements in the promoter regions of lettuce *LsTCP* genes.

