## Supplementary Figs 1-2 and Supplementary Tables 1-3 for "miR319 promotes *de novo* shoot regeneration by repressing *LsTCP4* in lettuce"

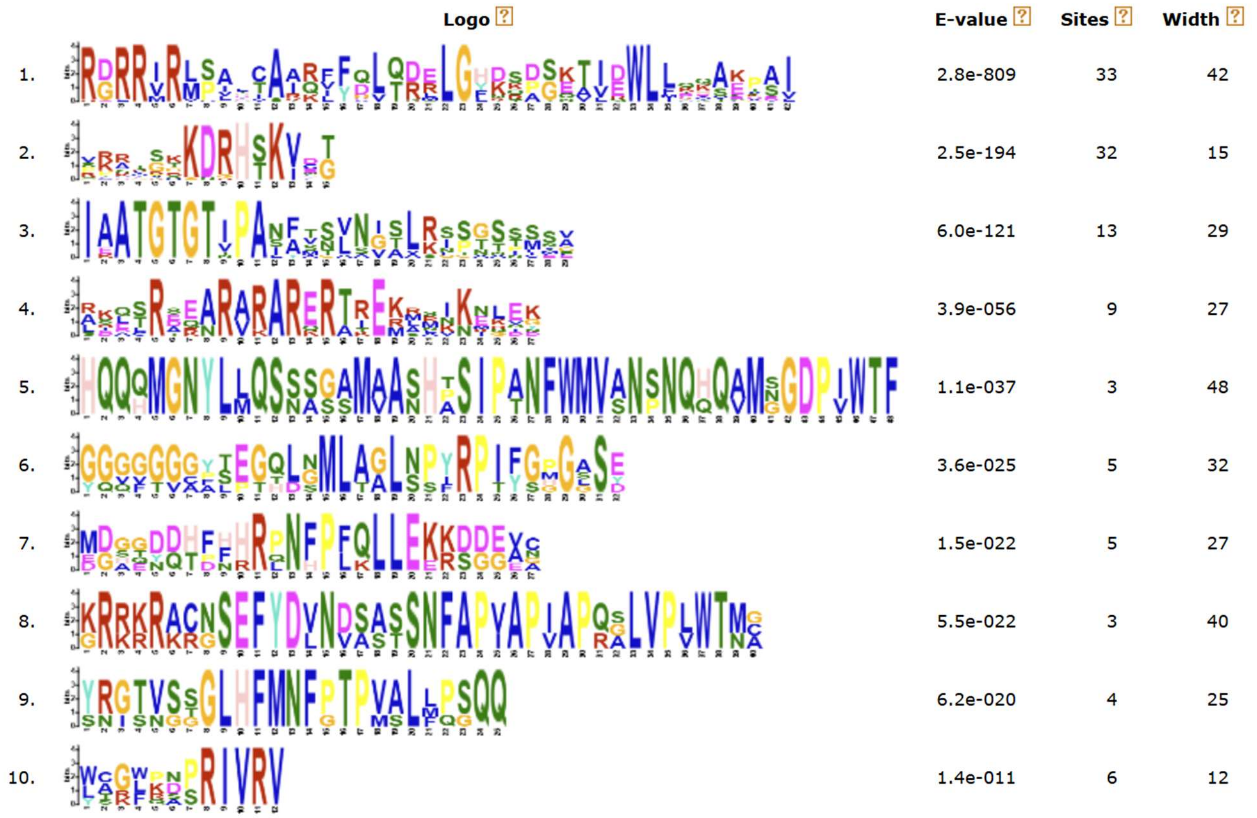


**Supplementary Fig. 1. Conserved motif sequence logos of lettuce LsTCP proteins.** Sequence logos show amino acid conservation patterns for the 10 motifs identified among LsTCP proteins using MEME Suite. Each logo represents the relative amino acid frequency at each position within the corresponding motif, with letter height indicating the degree of conservation. The E-value, number of motif sites, and motif width are shown at the right of each logo. Motif 1 was detected in all 33 LsTCP proteins, indicating that it represents a broadly conserved sequence feature within the family. The relationship between conserved motifs and the annotated TCP domain is shown in Fig. 1C.

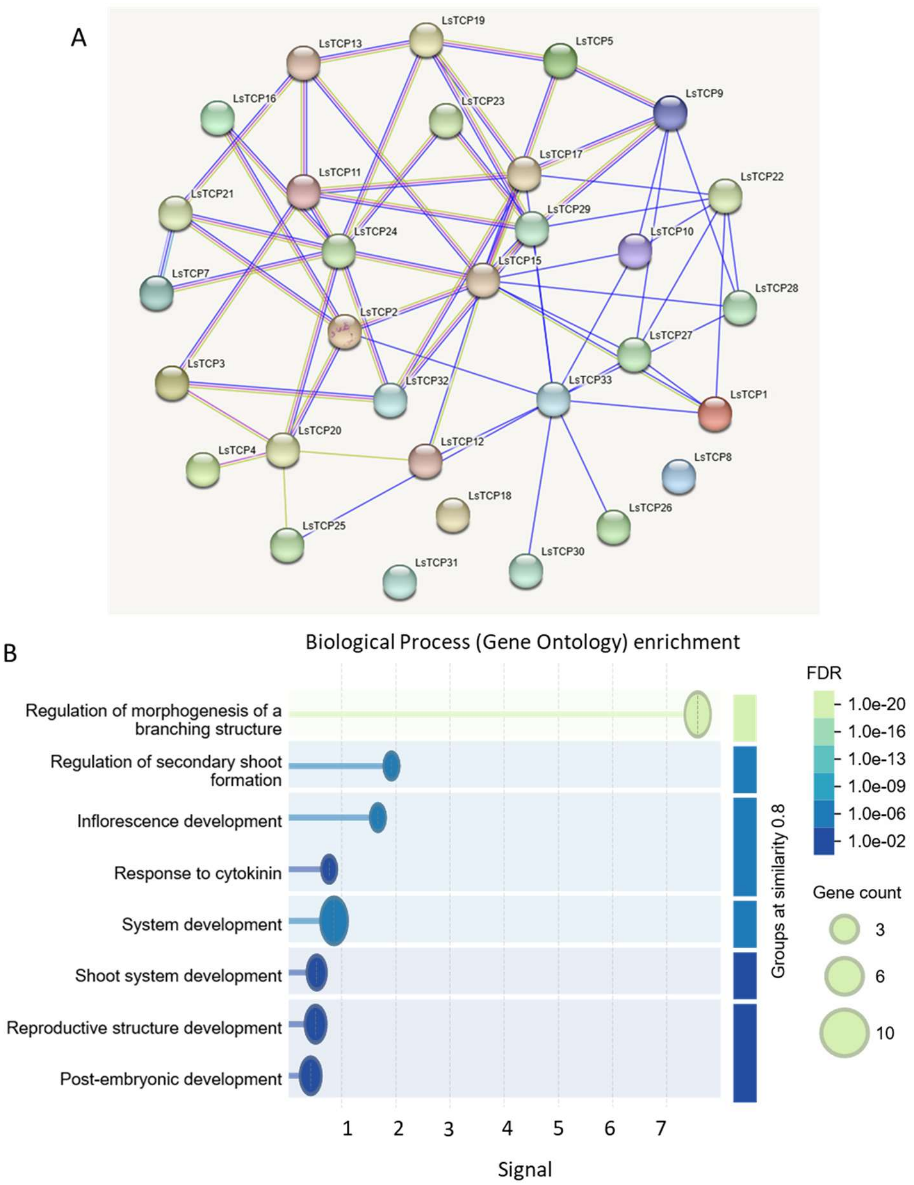


**Supplementary Fig. 2. Predicted protein-association network and Gene Ontology enrichment analysis of lettuce LsTCP proteins.** A, Predicted protein-association network of LsTCP proteins generated using STRING. Nodes represent individual LsTCP proteins, and edges represent predicted functional associations. B, Gene Ontology biological-process enrichment analysis of proteins represented in the predicted network, performed using ShinyGO. The bubble plot displays enriched developmental terms, including regulation of morphogenesis of a branching structure, regulation of secondary shoot formation, inflorescence development, response to cytokinin, system development, shoot-system development, reproductive-structure development, and post-embryonic development. The x-axis indicates enrichment signal, bubble size represents the number of genes assigned to each term, and bubble color indicates the false discovery rate (FDR).

**Supplementary Table 1.** Primers used for qRT-PCR analysis of selected *LsTCP* genes.

| **Gene name** | **Forward primer sequence （5'→3'）** | **Reverse primer sequence（5'→3'）** |
|---|---|---|
| *LsUBQ5* | AGAAGGAATCCACTCTCCACCTT | GCTTGGTGTAGGTCTTCTTCTTACG |
| *LsTCP2* | GGCGGCATCTACTTCCATTG | ACTGTTTCCCGTCGACTTCT |
| *LsTCP3* | TACGGTCAACTGGTCGGAAA | GAATTGGATGGCAGTGTGGG |
| *LsTCP4* | ACCCCTTCAGTCCAGTAACG | AGGGTTGGGTGTTGATCGAT |
| *LsTCP10* | AAGGAGGTGGTCGAATCGTT | AGCTGATAAACGAACACGCC |
| *LsTCP13* | GAGGGTTAAGAGATCGGCGA | GAGGAGGGAGTTCGTCGATT |
| *LsTCP17* | AGTTTCACGAGCCTTTGGTG | AAGTCCGAGGCGATCTTGAA |
| *LsTCP24* | ACGCCGAATTTGAAGATCCG | TCGAGCTCTGACCCGATTTT |
| *MIR319a* | *ATCCAAATACCGAGTCGCCA* | GGAGCTCCCTTCAGTCCAA |

**Supplementary Table 2**. Physicochemical characterization of lettuce LsTCP proteins.

| Protein Name | Length (aa) | MW (kDa) | Theoretical pI | Instability Index | Aliphatic Index | GRAVY | Pfam (PF03634) | Subcellular localization |
|---|---|---|---|---|---|---|---|---|
| LsTCP1 | 300 | 34.58 | 7.98 | 53.27 | 59.77 | -0.826 | Yes | Nuclear |
| LsTCP2 | 462 | 49.84 | 9.19 | 49.41 | 53.83 | -0.885 | Yes | Nuclear |
| LsTCP3 | 353 | 38.53 | 6.13 | 58.24 | 55.01 | -0.730 | Yes | Nuclear |
| LsTCP4 | 532 | 58.99 | 6.37 | 60.85 | 57.59 | -0.940 | Yes | Nuclear |
| LsTCP5 | 341 | 38.21 | 7.13 | 42.29 | 68.89 | -0.687 | Yes | Nuclear |
| LsTCP6 | 400 | 42.99 | 8.42 | 53.32 | 58.60 | -0.622 | Yes | Nuclear |
| LsTCP7 | 239 | 25.53 | 9.69 | 57.70 | 69.08 | -0.451 | Yes | Nuclear |
| LsTCP8 | 544 | 58.92 | 6.78 | 56.77 | 54.63 | -0.830 | Yes | Nuclear |
| LsTCP9 | 298 | 31.90 | 9.79 | 41.15 | 64.93 | -0.395 | Yes | Nuclear |
| LsTCP10 | 229 | 26.23 | 5.25 | 56.04 | 68.95 | -0.798 | Yes | Nuclear |
| LsTCP11 | 304 | 32.65 | 8.26 | 56.73 | 74.47 | -0.284 | Yes | Nuclear |
| LsTCP12 | 403 | 45.66 | 6.28 | 58.05 | 54.99 | -0.994 | Yes | Nuclear |
| LsTCP13 | 332 | 37.33 | 6.76 | 53.16 | 62.50 | -0.858 | Yes | Nuclear |
| LsTCP14 | 392 | 42.40 | 7.27 | 54.68 | 62.50 | -0.629 | Yes | Nuclear |
| LsTCP15 | 321 | 34.47 | 7.13 | 46.06 | 58.72 | -0.648 | Yes | Nuclear |
| LsTCP16 | 407 | 42.80 | 6.93 | 60.27 | 59.29 | -0.594 | Yes | Nuclear |
| LsTCP17 | 289 | 32.61 | 7.22 | 48.85 | 71.76 | -0.742 | Yes | Nuclear |
| LsTCP18 | 391 | 44.03 | 7.67 | 48.66 | 52.69 | -0.917 | Yes | Nuclear |
| LsTCP19 | 366 | 38.69 | 5.62 | 66.02 | 67.51 | -0.433 | Yes | Nuclear |
| LsTCP20 | 313 | 33.79 | 8.54 | 57.72 | 65.85 | -0.700 | Yes | Nuclear |
| LsTCP21 | 259 | 27.29 | 9.65 | 52.17 | 70.54 | -0.453 | Yes | Nuclear |
| LsTCP22 | 126 | 14.10 | 9.73 | 49.23 | 78.97 | -0.652 | Yes | Nuclear |
| LsTCP23 | 340 | 36.45 | 7.28 | 52.77 | 55.15 | -0.584 | Yes | Nuclear |
| LsTCP24 | 454 | 49.88 | 7.94 | 46.25 | 55.46 | -0.912 | Yes | Nuclear |
| LsTCP25 | 283 | 32.36 | 8.86 | 49.20 | 57.49 | -0.804 | Yes | Nuclear |
| LsTCP26 | 300 | 34.33 | 9.90 | 43.83 | 65.00 | -0.865 | Yes | Nuclear |
| LsTCP27 | 209 | 24.07 | 5.62 | 49.34 | 61.15 | -0.928 | Yes | Nuclear |
| LsTCP28 | 219 | 25.42 | 5.62 | 42.94 | 73.47 | -0.847 | Yes | Nuclear |
| LsTCP29 | 309 | 35.23 | 9.19 | 55.61 | 70.00 | -0.625 | Yes | Nuclear |
| LsTCP30 | 400 | 46.00 | 7.32 | 50.97 | 66.35 | -0.869 | Yes | Nuclear |
| LsTCP31 | 401 | 45.49 | 9.32 | 41.79 | 53.27 | -1.022 | Yes | Nuclear |
| LsTCP32 | 323 | 34.34 | 7.81 | 60.84 | 71.86 | -0.467 | Yes | Nuclear |
| LsTCP33 | 171 | 19.35 | 7.02 | 56.34 | 67.72 | -0.819 | Yes | Nuclear |

**Supplementary Table 3.** Ka/Ks analysis of selected *LsTCP* gene pairs.

| Paralog 1 | Paralog 2 | Ks | Ka | Ka/Ks | Selection pressure |
|---|---|---|---|---|---|
| *LsTCP1* | *LsTCP26* | 0.936 | 0.273 | 0.29202837 | Negative/Purifying |
| *LsTCP1* | *LsTCP4* | 1.830 | 0.652 | 0.3562724 | Negative/Purifying |
| *LsTCP1* | *LsTCP10* | 1.757 | 0.706 | 0.40193866 | Negative/Purifying |
| *LsTCP1* | *LsTCP25* | 0.853 | 0.340 | 0.39861089 | Negative/Purifying |
| *LsTCP2* | *LsTCP24* | 1.207 | 0.099 | 0.08175299 | Negative/Purifying |
| *LsTCP2* | *LsTCP27* | 3.167 | 0.667 | 0.21052262 | Negative/Purifying |
| *LsTCP26* | *LsTCP6* | 1.294 | 1.095 | 0.84622092 | Negative/Purifying |
| *LsTCP26* | *LsTCP10* | 2.009 | 0.606 | 0.30143421 | Negative/Purifying |
| *LsTCP26* | *LsTCP14* | 1.116 | 1.191 | 1.06704915 | Positive/Diversifying |
| *LsTCP26* | *LsTCP16* | 1.351 | 1.127 | 0.83423764 | Negative/Purifying |
| *LsTCP26* | *LsTCP18* | 2.024 | 0.461 | 0.22795306 | Negative/Purifying |
| *LsTCP26* | *LsTCP20* | 2.533 | 1.071 | 0.42280217 | Negative/Purifying |
| *LsTCP26* | *LsTCP25* | 1.154 | 0.335 | 0.29016431 | Negative/Purifying |
| *LsTCP26* | *LsTCP27* | 1.468 | 0.708 | 0.48254324 | Negative/Purifying |
| *LsTCP26* | *LsTCP28* | 1.989 | 0.624 | 0.3136551 | Negative/Purifying |
| *LsTCP4* | *LsTCP6* | 2.117 | 0.923 | 0.43610145 | Negative/Purifying |
| *LsTCP4* | *LsTCP9* | 2.005 | 0.988 | 0.49284173 | Negative/Purifying |
| *LsTCP4* | *LsTCP13* | 3.067 | 0.506 | 0.16504426 | Negative/Purifying |
| *LsTCP4* | *LsTCP14* | 3.044 | 0.907 | 0.29798033 | Negative/Purifying |
| *LsTCP4* | *LsTCP15* | 1.391 | 0.865 | 0.62199958 | Negative/Purifying |
| *LsTCP4* | *LsTCP16* | 1.620 | 0.919 | 0.5668464 | Negative/Purifying |
| *LsTCP4* | *LsTCP21* | 2.549 | 0.968 | 0.37960053 | Negative/Purifying |
| *LsTCP4* | *LsTCP22* | 1.560 | 1.057 | 0.6774191 | Negative/Purifying |
| *LsTCP4* | *LsTCP23* | 2.428 | 0.979 | 0.40342369 | Negative/Purifying |
| *LsTCP4* | *LsTCP24* | 2.871 | 0.464 | 0.16157368 | Negative/Purifying |
| *LsTCP4* | *LsTCP3* | 1.063 | 0.101 | 0.09488333 | Negative/Purifying |
| *LsTCP4* | *LsTCP27* | 2.781 | 0.385 | 0.13836763 | Negative/Purifying |
| *LsTCP4* | *LsTCP30* | 2.033 | 0.658 | 0.32345711 | Negative/Purifying |
| *LsTCP4* | *LsTCP32* | 1.782 | 1.141 | 0.64045354 | Negative/Purifying |
| *LsTCP4* | *LsTCP33* | 1.774 | 0.973 | 0.54822574 | Negative/Purifying |
| *LsTCP5* | *LsTCP10* | 2.056 | 0.489 | 0.23785388 | Negative/Purifying |
| *LsTCP5* | *LsTCP18* | 1.925 | 0.726 | 0.37714584 | Negative/Purifying |
| *LsTCP5* | *LsTCP27* | 3.072 | 0.583 | 0.18989584 | Negative/Purifying |
| *LsTCP5* | *LsTCP29* | 1.016 | 0.297 | 0.29270067 | Negative/Purifying |
| *LsTCP5* | *LsTCP31* | 1.877 | 0.774 | 0.41218311 | Negative/Purifying |
| *LsTCP6* | *LsTCP7* | 2.228 | 0.363 | 0.16276361 | Negative/Purifying |
| *LsTCP6* | *LsTCP10* | 2.737 | 0.946 | 0.3454773 | Negative/Purifying |
| *LsTCP6* | *LsTCP12* | 3.125 | 0.960 | 0.30726481 | Negative/Purifying |
| *LsTCP6* | *LsTCP13* | 2.382 | 1.094 | 0.45934836 | Negative/Purifying |
| *LsTCP6* | *LsTCP14* | 1.036 | 0.122 | 0.11761282 | Negative/Purifying |
| *LsTCP6* | *LsTCP15* | 2.413 | 0.303 | 0.12556894 | Negative/Purifying |
| *LsTCP6* | *LsTCP16* | 1.048 | 0.069 | 0.06627474 | Negative/Purifying |

| | | | | | |
|---|---|---|---|---|---|
| *LsTCP6* | *LsTCP18* | 1.994 | 0.979 | 0.49077284 | Negative/Purifying |
| *LsTCP6* | *LsTCP25* | 2.868 | 1.062 | 0.37009676 | Negative/Purifying |
| *LsTCP6* | *LsTCP27* | 1.589 | 1.102 | 0.69392125 | Negative/Purifying |
| *LsTCP7* | *LsTCP8* | 2.395 | 0.395 | 0.16486055 | Negative/Purifying |
| *LsTCP7* | *LsTCP16* | 1.729 | 0.385 | 0.22276815 | Negative/Purifying |
| *LsTCP7* | *LsTCP18* | 3.213 | 1.016 | 0.31623822 | Negative/Purifying |
| *LsTCP7* | *LsTCP21* | 1.902 | 0.110 | 0.05781626 | Negative/Purifying |
| *LsTCP7* | *LsTCP23* | 2.377 | 0.421 | 0.17695654 | Negative/Purifying |
| *LsTCP7* | *LsTCP24* | 2.410 | 1.092 | 0.4532023 | Negative/Purifying |
| *LsTCP8* | *LsTCP10* | 3.007 | 0.874 | 0.29079763 | Negative/Purifying |
| *LsTCP8* | *LsTCP23* | 1.883 | 0.330 | 0.17517673 | Negative/Purifying |
| *LsTCP8* | *LsTCP24* | 2.698 | 1.048 | 0.38836055 | Negative/Purifying |
| *LsTCP8* | *LsTCP27* | 1.256 | 1.173 | 0.93437685 | Negative/Purifying |
| *LsTCP9* | *LsTCP25* | 2.253 | 1.067 | 0.47376361 | Negative/Purifying |
| *LsTCP9* | *LsTCP3* | 2.090 | 0.941 | 0.45033136 | Negative/Purifying |
| *LsTCP10* | *LsTCP12* | 3.199 | 0.693 | 0.21647605 | Negative/Purifying |
| *LsTCP10* | *LsTCP13* | 2.214 | 0.507 | 0.2288754 | Negative/Purifying |
| *LsTCP10* | *LsTCP25* | 1.477 | 0.634 | 0.42928853 | Negative/Purifying |
| *LsTCP10* | *LsTCP27* | 2.846 | 0.400 | 0.14044225 | Negative/Purifying |
| *LsTCP11* | *LsTCP17* | 2.090 | 1.069 | 0.51147692 | Negative/Purifying |
| *LsTCP11* | *LsTCP19* | 0.925 | 0.122 | 0.13149205 | Negative/Purifying |
| *LsTCP11* | *LsTCP24* | 3.036 | 0.995 | 0.32777085 | Negative/Purifying |
| *LsTCP11* | *LsTCP25* | 2.584 | 1.010 | 0.3907451 | Negative/Purifying |
| *LsTCP11* | *LsTCP30* | 1.736 | 1.133 | 0.65254476 | Negative/Purifying |
| *LsTCP11* | *LsTCP31* | 1.983 | 1.021 | 0.51485344 | Negative/Purifying |
| *LsTCP11* | *LsTCP32* | 0.616 | 0.135 | 0.2191429 | Negative/Purifying |
| *LsTCP12* | *LsTCP13* | 1.398 | 0.617 | 0.44141959 | Negative/Purifying |
| *LsTCP12* | *LsTCP16* | 1.804 | 0.914 | 0.5063047 | Negative/Purifying |
| *LsTCP12* | *LsTCP17* | 1.823 | 0.653 | 0.35835634 | Negative/Purifying |
| *LsTCP12* | *LsTCP20* | 2.931 | 1.008 | 0.34374723 | Negative/Purifying |
| *LsTCP12* | *LsTCP27* | 1.579 | 0.756 | 0.47888417 | Negative/Purifying |
| *LsTCP12* | *LsTCP28* | 2.947 | 0.677 | 0.22984905 | Negative/Purifying |
| *LsTCP12* | *LsTCP30* | 1.322 | 0.218 | 0.16454804 | Negative/Purifying |
| *LsTCP13* | *LsTCP14* | 1.934 | 1.043 | 0.53958925 | Negative/Purifying |
| *LsTCP13* | *LsTCP16* | 2.955 | 1.035 | 0.35008961 | Negative/Purifying |
| *LsTCP13* | *LsTCP17* | 1.503 | 0.163 | 0.10822363 | Negative/Purifying |
| *LsTCP13* | *LsTCP27* | 1.830 | 0.577 | 0.31525445 | Negative/Purifying |
| *LsTCP13* | *LsTCP28* | 1.432 | 0.493 | 0.34394602 | Negative/Purifying |
| *LsTCP14* | *LsTCP16* | 1.130 | 0.075 | 0.06630332 | Negative/Purifying |
| *LsTCP14* | *LsTCP18* | 2.945 | 0.905 | 0.3072916 | Negative/Purifying |
| *LsTCP14* | *LsTCP22* | 2.421 | 0.610 | 0.25194504 | Negative/Purifying |
| *LsTCP14* | *LsTCP25* | 2.864 | 1.023 | 0.35714988 | Negative/Purifying |
| *LsTCP14* | *LsTCP3* | 2.375 | 0.942 | 0.39648148 | Negative/Purifying |
| *LsTCP14* | *LsTCP28* | 2.126 | 1.154 | 0.54284334 | Negative/Purifying |
| *LsTCP14* | *LsTCP30* | 2.216 | 1.010 | 0.45597452 | Negative/Purifying |

| | | | | | |
|---|---|---|---|---|---|
| *LsTCP15* | *LsTCP16* | 1.455 | 0.320 | 0.21980024 | Negative/Purifying |
| *LsTCP15* | *LsTCP18* | 1.497 | 0.889 | 0.5938911 | Negative/Purifying |
| *LsTCP15* | *LsTCP21* | 1.821 | 0.381 | 0.20909546 | Negative/Purifying |
| *LsTCP15* | *LsTCP23* | 1.863 | 0.274 | 0.14699342 | Negative/Purifying |
| *LsTCP15* | *LsTCP3* | 1.344 | 0.792 | 0.58959473 | Negative/Purifying |
| *LsTCP15* | *LsTCP30* | 3.198 | 0.996 | 0.31149512 | Negative/Purifying |
| *LsTCP15* | *LsTCP31* | 2.585 | 0.821 | 0.31755944 | Negative/Purifying |
| *LsTCP15* | *LsTCP32* | 2.598 | 0.404 | 0.15563495 | Negative/Purifying |
| *LsTCP15* | *LsTCP33* | 2.179 | 0.514 | 0.23581582 | Negative/Purifying |
| *LsTCP16* | *LsTCP18* | 2.516 | 0.935 | 0.37154708 | Negative/Purifying |
| *LsTCP16* | *LsTCP20* | 2.074 | 0.404 | 0.19474683 | Negative/Purifying |
| *LsTCP16* | *LsTCP21* | 1.461 | 0.364 | 0.24897385 | Negative/Purifying |
| *LsTCP16* | *LsTCP23* | 2.130 | 0.160 | 0.07530956 | Negative/Purifying |
| *LsTCP16* | *LsTCP27* | 2.811 | 1.033 | 0.36758329 | Negative/Purifying |
| *LsTCP16* | *LsTCP28* | 1.490 | 1.144 | 0.76765197 | Negative/Purifying |
| *LsTCP16* | *LsTCP30* | 2.441 | 1.020 | 0.41761998 | Negative/Purifying |
| *LsTCP17* | *LsTCP19* | 3.015 | 1.013 | 0.33586806 | Negative/Purifying |
| *LsTCP17* | *LsTCP27* | 1.454 | 0.648 | 0.44578209 | Negative/Purifying |
| *LsTCP17* | *LsTCP28* | 2.448 | 0.504 | 0.20574588 | Negative/Purifying |
| *LsTCP17* | *LsTCP29* | 1.639 | 0.291 | 0.17739089 | Negative/Purifying |
| *LsTCP17* | *LsTCP30* | 1.195 | 0.639 | 0.5350644 | Negative/Purifying |
| *LsTCP17* | *LsTCP32* | 1.364 | 1.021 | 0.74910019 | Negative/Purifying |
| *LsTCP18* | *LsTCP27* | 1.286 | 0.772 | 0.59979958 | Negative/Purifying |
| *LsTCP18* | *LsTCP28* | 1.453 | 0.765 | 0.52633506 | Negative/Purifying |
| *LsTCP18* | *LsTCP31* | 1.492 | 0.205 | 0.13764613 | Negative/Purifying |
| *LsTCP18* | *LsTCP33* | 2.476 | 1.233 | 0.49779629 | Negative/Purifying |
| *LsTCP19* | *LsTCP22* | 3.176 | 0.564 | 0.17775868 | Negative/Purifying |
| *LsTCP19* | *LsTCP25* | 2.033 | 0.966 | 0.47521774 | Negative/Purifying |
| *LsTCP19* | *LsTCP27* | 2.186 | 1.224 | 0.55984532 | Negative/Purifying |
| *LsTCP19* | *LsTCP30* | 1.959 | 1.069 | 0.5458444 | Negative/Purifying |
| *LsTCP19* | *LsTCP32* | 0.647 | 0.086 | 0.13221898 | Negative/Purifying |
| *LsTCP19* | *LsTCP33* | 3.128 | 0.520 | 0.16625971 | Negative/Purifying |
| *LsTCP22* | *LsTCP23* | 2.630 | 0.581 | 0.22098151 | Negative/Purifying |
| *LsTCP22* | *LsTCP3* | 2.261 | 1.104 | 0.48847517 | Negative/Purifying |
| *LsTCP22* | *LsTCP33* | 2.417 | 0.374 | 0.15485406 | Negative/Purifying |
| *LsTCP23* | *LsTCP24* | 2.130 | 1.015 | 0.47662013 | Negative/Purifying |
| *LsTCP23* | *LsTCP27* | 2.194 | 0.926 | 0.42216339 | Negative/Purifying |
| *LsTCP23* | *LsTCP31* | 2.356 | 0.861 | 0.36549191 | Negative/Purifying |
| *LsTCP3* | *LsTCP28* | 2.144 | 0.345 | 0.16078609 | Negative/Purifying |
| *LsTCP3* | *LsTCP33* | 1.972 | 1.027 | 0.5208194 | Negative/Purifying |
| *LsTCP27* | *LsTCP28* | 1.151 | 0.280 | 0.24308325 | Negative/Purifying |
| *LsTCP27* | *LsTCP29* | 1.631 | 0.708 | 0.43401969 | Negative/Purifying |
| *LsTCP27* | *LsTCP32* | 2.148 | 1.256 | 0.58473029 | Negative/Purifying |
| *LsTCP27* | *LsTCP33* | 3.093 | 1.257 | 0.40645416 | Negative/Purifying |
| *LsTCP28* | *LsTCP29* | 2.001 | 0.603 | 0.3012604 | Negative/Purifying |

| | | | | | |
|---|---|---|---|---|---|
| *LsTCP28* | *LsTCP30* | 2.197 | 0.636 | 0.28962367 | Negative/Purifying |
| *LsTCP28* | *LsTCP33* | 2.478 | 1.312 | 0.52943335 | Negative/Purifying |
| *LsTCP29* | *LsTCP31* | 2.532 | 0.851 | 0.3360837 | Negative/Purifying |
| *LsTCP30* | *LsTCP32* | 1.287 | 1.123 | 0.87225173 | Negative/Purifying |

Note: Ka, nonsynonymous substitution rate; Ks, synonymous substitution rate. Ka/Ks < 1, = 1, and > 1 indicate patterns consistent with purifying, neutral, and diversifying selection, respectively.
